# A dynamic 3D polymer model of the *Escherichia coli* chromosome driven by data from optical pooled screening

**DOI:** 10.1101/2024.10.30.621082

**Authors:** Dvir Schirman, Konrad Gras, Vinodh Kandavalli, Jimmy Larsson, David Fange, Johan Elf

## Abstract

The DNA of bacterial cells is organized in a highly dynamic chromosome structure. To guarantee its propagation, the chromosome must replicate, segregate, and accommodate other biological processes like gene expression. Therefore, to understand the causal relationship between chromosome organization and biological function, it is essential to follow chromosome dynamics. Live-cell imaging of fluorescent labels allows the tracking of specific genomic locations, but the current imaging-based approaches to study bacterial chromosome organization over the cell cycle are limited in their throughput. Even with multicolor fluorescent locus labeling, the genomic resolution is insufficient to gain insights about the whole chromosome structure from a single experiment. Although sequencing-based methods like chromosome conformation capture provide high genomic coverage, they can only be done in bulk, providing an averaged static view of the chromosome structure. In this study, we address these limitations and investigate the bacterial chromosome organization by imaging fluorescently labeled loci in a library of *Escherichia coli* strains, following the 3D locations of 68 different loci in live cells in a single experiment. The resulting location distributions along the cell’s longitudinal and radial axes are used to inform a dynamic polymer model of the chromosome over the cell cycle. We show a global reorganization of the *E. coli* chromosome, at both the longitudinal and radial axes. Our model reproduces the known chromosomal architecture of four macrodomains and demonstrates how these domains form dynamically over the cell cycle.

## Introduction

Chromosome organization in *Escherichia coli* has been studied using various methods. A number of efforts have relied on DNA fluorescence in situ hybridization (FISH) to different chromosomal loci to elucidate their locations in fixed cells (Niki, Yamaichi, and Hiraga 2000; Bates and Kleckner 2005). These studies suggest that, at the Mbp scale, the chromosome in *E. coli* is organized into macrodomains corresponding to the origin (*oriC*) and terminus of replication (*ter*), with each domain being composed of loci that preferentially colocalize (Niki, Yamaichi, and Hiraga 2000). Additionally, studies based on FISH indicated that different loci occupy specific intracellular regions (Bates and Kleckner 2005). The use of DNA FISH has also been extended to perform highly multiplexed experiments for the study of chromosome organization in eukaryotic cells (Bintu et al. 2018). Despite this possibility for multiplexing, DNA FISH requires the fixation of cells and does not provide information about the dynamics of the chromosome over the division cycle.

Another set of methods that rely on fluorescence imaging is based on the specific binding between a fluorescently labeled DNA-binding protein to a DNA sequence introduced on the chromosome. These include different fluorescent repressor-operator systems (FROS) (Lau et al. 2004) and the ParB/*parS* system (Wang et al. 2006; Nielsen et al. 2006; Espeli, Mercier, and Boccard 2008), which have been used to label specific chromosomal loci and determine their intracellular locations. As this locus labeling approach allows for live-cell imaging it has also been used for tracking loci at short-time scales (Javer et al. 2013, 2014) and over the division cycle of the cells (Lau et al. 2004; Wang et al. 2006; Nielsen et al. 2006; Espéli et al. 2012; Youngren et al. 2014; Cass et al. 2016). Using this method, loci near *oriC* and *ter* have been found to localize towards the cell poles (Youngren et al. 2014), while the left and right chromosome arms occupy different cell halves (Wang et al. 2006; Nielsen et al. 2006) or align between the cell poles (Youngren et al. 2014). The different localization patterns for the left and right chromosome arms have been described as growth rate-dependent (Youngren et al. 2014; Badrinarayanan, Le, and Laub 2015), although a recent study suggests that both modes of organization exist across different growth conditions (Sadhir and Murray 2023). Additionally, several loci have been shown to occupy specific regions along the cell’s short axis (Youngren et al. 2014). While single-locus labeling provides a live-cell method to study locus localization, it suffers from low throughput, as only a limited number of loci can be studied simultaneously in each experiment.

Imaging of fluorescently labeled nucleoid-associated proteins (NAPs) has been employed to study the dynamics of the whole nucleoid (Fisher et al. 2013). Several of these proteins, such as HU and Fis, are abundant and bind throughout the chromosome, making it possible to use them as reporters for the global chromosome behavior in live cells. However, nucleoid imaging lacks the localization information for specific loci provided by DNA FISH and locus labeling with fluorescent DNA-binding proteins.

In contrast to fluorescence microscopy-based methods, chromosome conformation capture (3C) relies on measuring contacts between different loci to study bacterial chromosome organization. Initially developed to investigate eukaryotic chromosome organization (Dekker et al. 2002), this set of methods has also been applied to study bacterial chromosome biology (Le et al. 2013; Lioy et al. 2018; Marbouty et al. 2015). Applications of Hi-C, a chromosome conformation capture method that uses next-generation sequencing for quantifying pairwise locus contacts, have corroborated the presence of macrodomains (Lioy et al. 2018) and also identified chromosome interaction domains (CIDs) at the 30-400 kb scale (Le et al. 2013; Marbouty et al. 2015; Lioy et al. 2018). These structural domains appear to be delimited by regions of high gene expression (Lioy et al. 2018), although not consistently. Furthermore, recent applications of high-resolution Hi-C (∼5-10 kb) indicate regions of high transcriptional activity as a major constraining factor in *E. coli* chromosome organization (Bignaud et al. 2024). The first application of Hi-C was based on a bulk cell culture, but later developments have led to the method being used on single eukaryotic cells synchronized to the same cell cycle stage (Nagano et al. 2017; Ramani et al. 2017). However, a similar application to bacteria would be challenging, in part due to the lack of easily separable cell cycle stages in sequencing-based experiments. Thus, current Hi-C methods for *E. coli* are limited to bulk measurements that provide a population-averaged view of the chromosome.

Since the emergence of high-throughput 3C-based methods like Hi-C that capture chromosomal interactions at a whole genome level, there have been efforts to develop computational models to reconstruct the 3D chromosome structure based on interaction frequencies (Oluwadare, Highsmith, and Cheng 2019). A common approach to model the chromosome is by using a coarse-grained beads-on-a-string polymer (Mirny 2011). Most models do not use Hi-C-derived contact frequencies directly, but convert them to spatial distances by assuming an inverse power law relation between contact frequencies and pairwise distances (Fraser et al. 2009; Marbouty et al. 2015) or by calibrating contact frequencies with distances using imaging data (Umbarger et al. 2011). After converting a contact frequency matrix to an average distance matrix, a single consensus model that best fits the pairwise distances can be reconstructed (Umbarger et al. 2011; Marbouty et al. 2015; Hua and Ma 2019). However, unlike proteins which often have a specific structure, the chromosome structure is highly dynamic, therefore an expedient approach is to use molecular dynamics to reconstruct an ensemble of possible structures aiming to capture the population-level variability. In these approaches, distances are represented as spring-like interactions which constrain a dynamic model (Yildirim and Feig 2018; Wasim, Gupta, and Mondal 2021; Wasim, Bera, and Mondal 2023). A challenge in modeling bacterial chromosomes such as *E. coli* is that since they lack distinct cell cycle stages, the chromosome is continuously replicating and segregating as the cell grows and divides. Therefore, at any given stage of the replication cycle, the modeled polymer might have a different topology. Wasim et al. have tackled this challenge by combining discrete replication stages in their model (Wasim, Gupta, and Mondal 2021), and recent theoretical work has introduced a dynamic simulation of a replicating bacterial chromosome (Harju, van Teeseling, and Broedersz 2024).

To tackle the limitations of existing methods used to investigate the *E. coli* chromosome structure, we imaged a strain library with 83 different fluorescently labeled loci uniformly distributed throughout the chromosome. The cells were grown and imaged in a microfluidic mother-machine chip to determine the locus locations in live cells over their division cycle. We engineered the point-spread-function of the fluorescent labels and applied our neural network for three-dimensional (3D) localization of fluorescent foci (Karempudi et al. 2023). The resulting 3D locus location distributions were demultiplexed by genotyping of chromosomally expressed barcode sequences (Soares et al. 2023) in the same microfluidic chip. Finally, a 3D chromosome structure model was constructed by developing a polymer model informed by the demultiplexed locus location distributions along the cell’s long and radial axes.

## Results

### Mapping of pooled localization phenotypes to specific loci

We aimed to capture the dynamics of the three-dimensional (3D) chromosome over the cell cycle in living *Escherichia coli*. The only techniques that provide localization information in live cells use fluorescent DNA-binding proteins, e.g. FROS labeling (Lau et al. 2004). Thus, we used a single FROS-based label but introduced it at 83 different loci in separate *E. coli* strains that were pooled into a strain library (Fig. 1a). Pooling of the labeled strains made it possible to determine the locations of 83 different loci in a single microfluidics-based imaging experiment using optical pooled screening (Lawson et al. 2017). We introduced an array of twelve *malO* operators at specific loci on the chromosome. The *malO* array binds the maltose repressor MalI fused to the yellow fluorescent SYFP2, resulting in fluorescent foci that can be localized in live cells (Gras, Fange, and Elf 2024). However, localization of fluorescent emitters with a conventional wide-field fluorescence microscope cannot provide their 3D coordinates, only the coordinates in x and y. To acquire the emitter’s z-coordinates, we introduced astigmatism in the microscope with a cylindrical lens, which encodes the lateral information into the point-spread-function (PSF) of the fluorescent locus labels (Kao and Verkman 1994).

**Fig. 1:**
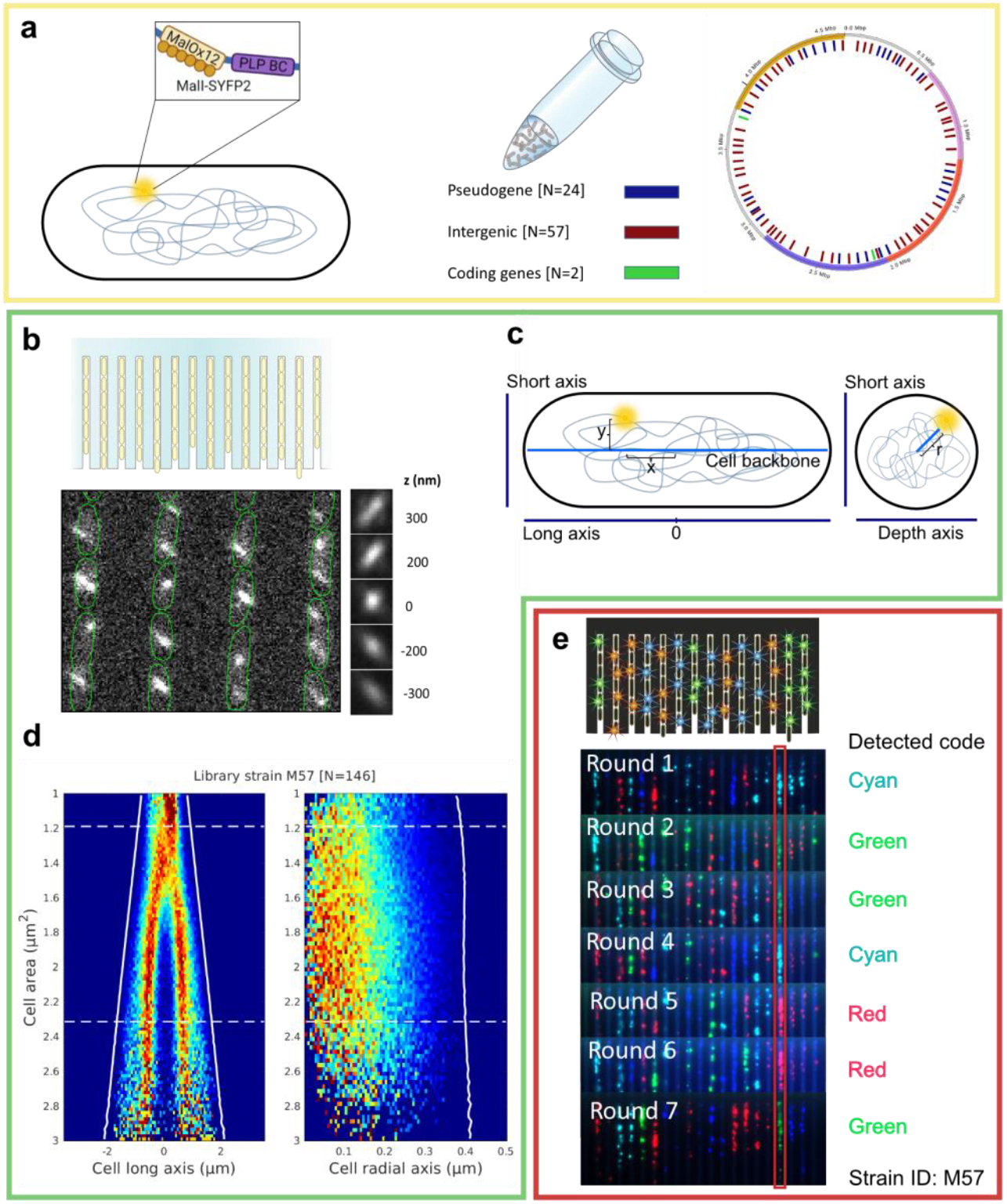
Phenotype-to-genotype mapping workflow. **a**, Cartoon illustrating the loci labeling and barcoding of a single *E. coli* strain (left). Chromosome map indicating where the loci labels of the strain library have been introduced on the chromosome (right). The lines showing the labels are color-coded based on the type of insertion site. The *oriC*, left, right and *ter* macrodomains are marked in yellow, purple, magenta and red based on (Espeli, Mercier, and Boccard 2008). **b**, An example fluorescence image of cells with fluorescent loci labels. The cell outlines (green) were determined using cell segmentation of phase contrast images. A stack of fluorescent beads imaged at selected z positions to demonstrate the astigmatic PSF. **c**, Cartoon illustrating the estimation of internal cell coordinates of the fluorescent emitters. The old cell pole is to the left in the cartoon and the new cell pole is to the right. **d**, Loci location distributions along the cell’s long and radial axes as a function of cell area. The white dashed lines indicate the average birth and division areas. The solid white lines indicate the average position of the cell poles, with the old cell pole corresponding to the left line and the new cell pole corresponding to the right line. Distributions are exemplified for a loci label 1 Mbp from *oriC* on the left replichore. The heat map colors are based on the number of detected fluorescent emitters per cell. **e**, Example fluorescence images from seven rounds of FISH used to determine the strain’s identity in the microfluidic sample. Fluorescence signals are based on hybridization between a fluorescent oligonucleotide and the expressed barcode RNA. The red box highlights the signals obtained from a single growth channel over all rounds to exemplify the genotyping analysis.

To study the chromosome structure over the bacterial cell cycle, we used a microfluidic mother-machine device to facilitate the continuous growth of the cells and image them while growing (Wang et al. 2010; Baltekin et al. 2017). We acquired phase contrast images every 2 minutes and fluorescence images every 4 minutes over 10 hours and assembled them into time lapses. Outlines of the cells were estimated by segmenting the phase contrast images (Fig. 1b) using the Omnipose network (Kandavalli et al. 2022; Cutler et al. 2022) and connected into tracks of cell lineages (Magnusson et al. 2015). The 3D locations of the fluorescent foci were determined (Fig. 1b) with our field-dependent DeepLoc neural network algorithm (Karempudi et al. 2023).

To visualize the 3D locus locations as a function of the cell cycle, we transformed the emitter image coordinates into the cell’s internal coordinate system (Fig. 1c). While the x and y coordinates can be transformed based on the length, width, and curvature of the cell outlines, there is no reference for the z-coordinate along the cell’s depth. The z-coordinate output from the neural network is relative to the PSF model, and hence we used the average of the z-coordinate distribution over the image plane at each position of the microfluidic chip to define zero along the depth axis (Karempudi et al. 2023). The internal coordinates were binned as a function of the area enclosed by the cell outlines (hereafter termed cell area) and visualized as bi-variate location distributions (Fig. 1d). The *E. coli* cell shape is radially symmetric, meaning that the only difference between the cell’s short and depth axes is rotation. Thus, instead of visualizing these axes separately, we plotted the normalized radial coordinates (hereafter termed radial coordinates) as a function of the cell area (Fig. 1d, see Materials & Methods for details on the normalization).

To allow for genotyping of each strain in the library, all strains had a unique chromosomal barcode sequence introduced adjacent to the *malO* array. Following the collection of the 3D localization data, a second experiment was performed in the same microfluidic chip to correlate genotype (i.e. the label’s location in the genome) and localization phenotype (Fig. 1e), as outlined in (Soares et al. 2023). Briefly, the cells were fixed and permeabilized with 70% EtOH, and the barcode sequences were expressed from the chromosome using T7 polymerase in the fixed cells (Askary et al. 2020), and amplified with rolling circle amplification. Genotyping was then performed with FISH by flowing in fluorescent detection probes that hybridize to the barcodes and detecting the resulting fluorescent signals in each channel. Strains were identified by performing seven sequential rounds of FISH probe hybridization, assigning a fluorescent color per strain in each round. The strain identity in each growth channel was determined by the unique sequence of fluorescent signals in that channel. This allowed us to map the labeled loci to a 3D localization phenotype (Fig. 1e). See Materials & Methods for details on the phenotype-to-genotype mapping.

### Capturing the intracellular locations of multiple loci over the cell cycle

Using our phenotype-to-genotype mapping approach, we determined the cell cycle-dependent location distributions along the cell’s long and radial axes for 68 out of the 83 loci in the library pooled from 4 independent repeats of the experiment. To check if the genotyping results faithfully represents the strains’ distribution in the pooled library, we performed amplicon deep sequencing and compared it with the genotyping (Fig. S1). In total, 15 strains were underrepresented in the genotyping (<10 channels), with 14 of these also being underrepresented in the pool based on sequencing, and hence were discarded from the analysis (Supplemental Information). The genotyping statistics for the pooled experiment are shown in Table S1.

We exemplify the distributions of eight selected loci in (Fig. 2). The exemplified long axis localization patterns have been observed previously (Youngren et al. 2014; Wallden et al. 2016; Knöppel et al. 2023; Sadhir and Murray 2023; Gras, Fange, and Elf 2024), with *oriC*-adjacent loci being found near the old cell pole early in the cell cycle followed by movement of the replicated locus copies towards the quarter long axis positions. In our experimental growth conditions the cells have an average generation time of 60 minutes, where the cells are born with an ongoing round of replication and two copies of *oriC* (Wallden et al. 2016; Gras, Fange, and Elf 2024). Localization distributions of all labeled loci in the library are shown in Fig. S2. Thus, our approach successfully reproduces previously shown long axis locations of loci over the cell cycle, and with fewer experiments than if each labeled locus would have been imaged separately.

**Fig. 2:**
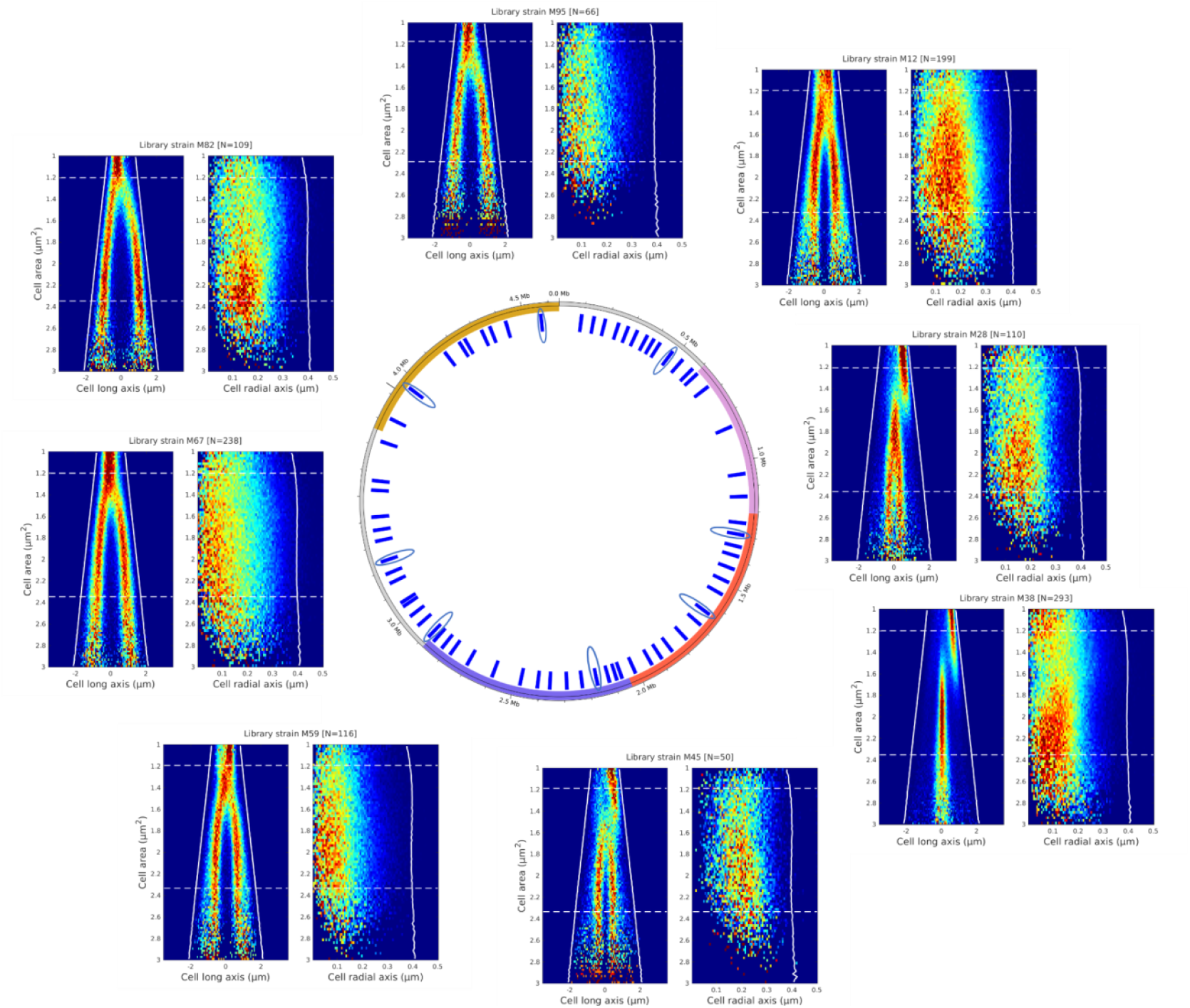
Examples of demultiplexed location distributions over the cell cycle. Loci intracellular location distributions along the cell’s long axis and radial axis as a function of cell area. Distributions of eight selected loci from the strain library are shown. Long axis and radial axis location distributions as in Fig. 1d. The tall tick in the chromosome map indicates *oriC*.

Some of the radial localization patterns have been observed indirectly by examining locus localization distributions in the yz-plane (Karempudi et al. 2023) or inferred from short axis distributions (Youngren et al. 2014; Gras, Fange, and Elf 2024). Although the radial location distributions for each loci remained relatively constant throughout the cell cycle (Fig. 2), the average radial positioning differed between them, showing that the loci occupy distinct radial regions in the cell.

### Global dynamics of the chromosome structure

To get an overview of all loci positions inside the cells, we assembled the demultiplexed localization data from all labeled loci and plotted it as a function of the chromosome sequence coordinate for a set of different cell areas. Notably, the loci’s absolute internal long axis coordinates (Fig. 3) showed a movement towards the middle of the cell as the cell cycle progressed, likely due to replication (Gras, Fange, and Elf 2024), followed by relocation of non-*ter* loci away from the middle. Loci near *ter* (Fig. 3, genomic loci around 1.5 - 2 Mbp) remained at the cell middle late into the cell cycle, while loci outside of *ter* extended almost linearly towards *oriC* at the quarter-long axis positions. To examine the dynamics of the whole chromosome longitudinally, we plotted the loci positions as in Fig. 3, but for 30 cell area bins and assembled them into Movie S1.

**Fig. 3:**
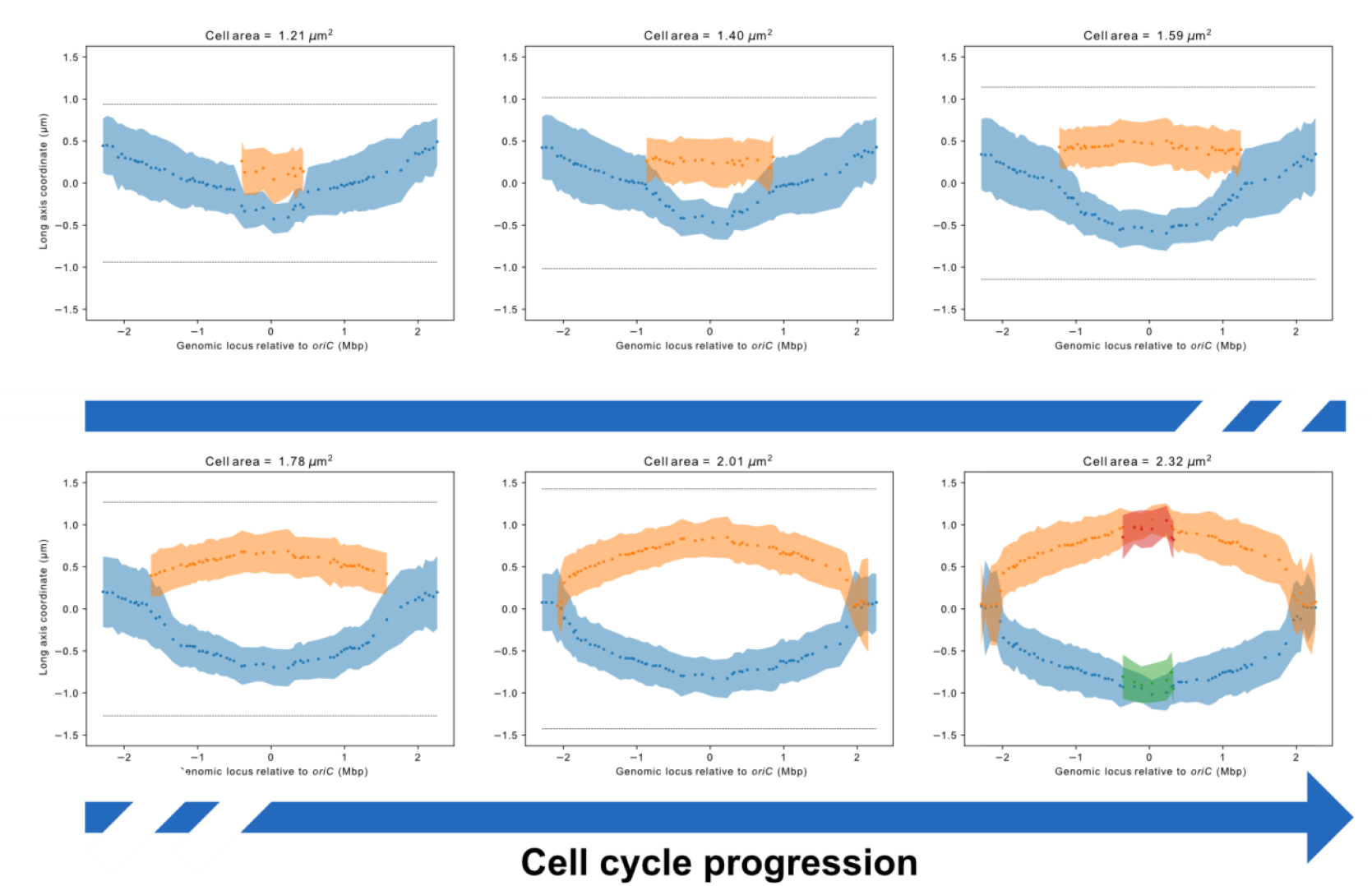
Locus long axis locations over the cell cycle. The average loci long axis coordinates as a function of the chromosome sequence position at selected cell areas. Each point corresponds to the average long axis coordinate based on a fit of the long axis coordinate distributions in Fig. S1. Shaded areas show standard deviation of the fit to the location distributions. The dotted lines show the average positions of the old and new cell poles, with the new pole at the top and old pole at the bottom. Points in blue are based on the location distribution branch closest to the old cell pole and the orange points are based on branches closest to the new cell pole. Green and purple points are based on the branches that segregate away from the old and new cell poles, respectively.

To examine the radial dynamics of the chromosome, we assembled the radial localization data over the cell cycle (Movie S2), but for most loci the radial positioning was relatively constant throughout the cell cycle. Visualizing the radial coordinates together in an averaged plot of Movie S2 revealed regions of several kb that exhibited distinct radial localization patterns (Fig. 4a). This was most clear for loci in the *ter*-region, which were radially localized near the cell center, and the regions on the left and right of *ter*, which were found further away from the center. The seemingly symmetric pattern centered around *ter* led us to investigate the average radial coordinates as a function of genomic distance from *oriC* (Fig. 4b). Indeed, loci 1.2-2.06 Mbp away from *oriC* on both chromosome arms showed the largest mean radial coordinates. We also observed a 74% and 65% overlap (Jaccard index) between the outwardly located regions and the left and right macrodomains respectively (Fig. 4a).

**Fig. 4:**
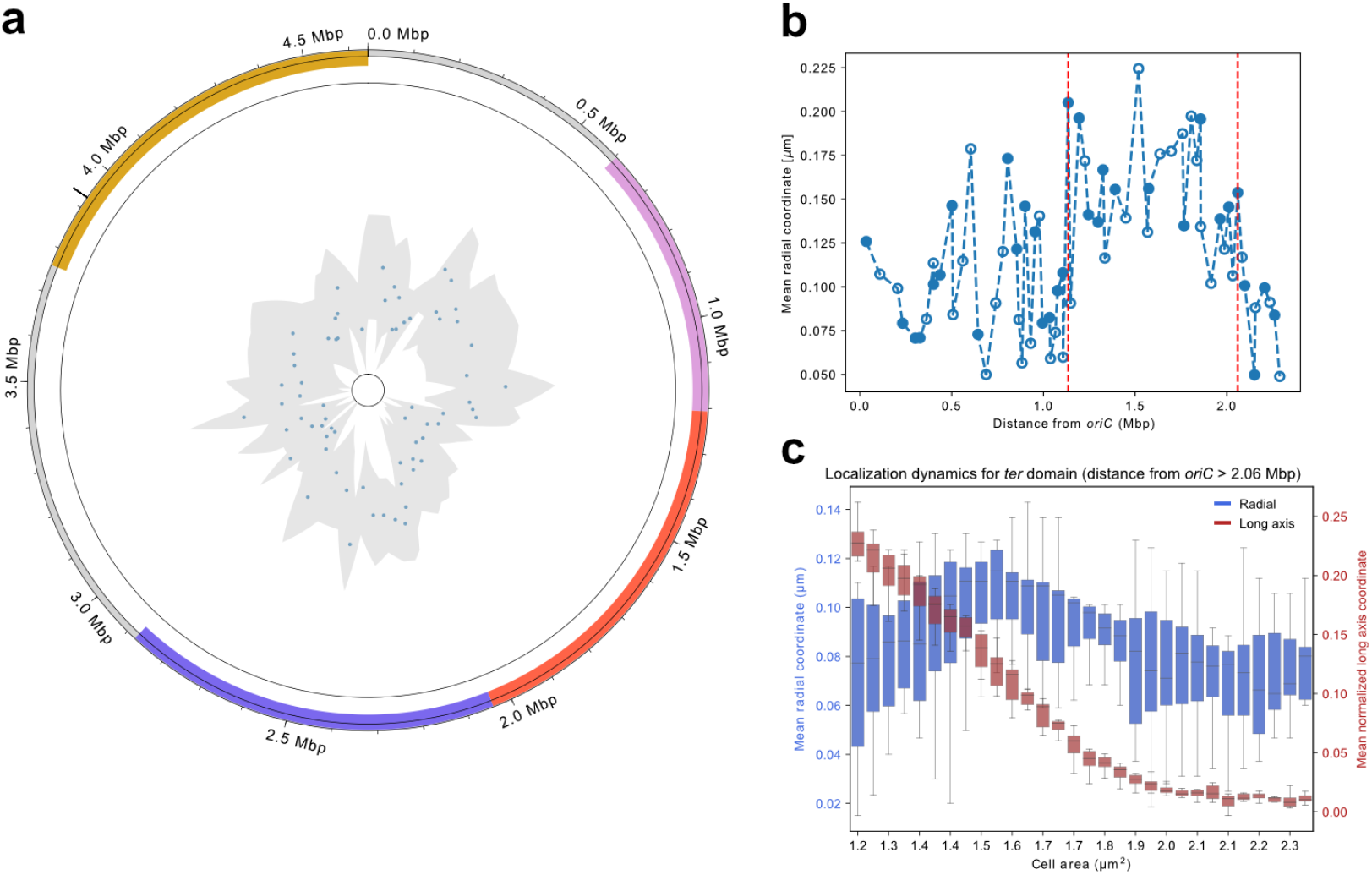
Radial locus locations across the chromosome. **a**, Radial distribution coordinates for each locus label in the strain library over the chromosome. Points correspond to the mode of the radial location distribution of each locus label, shaded areas correspond to the standard deviation. Mode and standard deviation are calculated from the empirical distribution of radial coordinates after normalizing by the area associated with each radial bin. The central circle corresponds to 0 along the cell’s radial axis and the outer circle corresponds to the cell membrane (r = 0.47 μm). The *oriC*, left, right and *ter* macrodomains are indicated in yellow, blue, pink and red respectively, based on (Espeli, Mercier, and Boccard 2008). **b**, Mean radial loci coordinates as a function of distance from *oriC* averaged over all cell areas in the corresponding radial axis distributions (Fig. S2). Red dashed lines indicate loci within 1.2-2.06 Mbp from *oriC*. Empty circles represent loci to the left of *oriC*, and full circles are loci to the right of *oriC*. **c**, Repositioning of the *ter* loci (>2.06 Mbp from *oriC*) along the cell’s radial and long axes as a function of cell area. The box plots are based on combined mean coordinates for the selected loci. Box edges represent the quartiles of the distribution, whiskers represent the extreme values.

The average radial coordinates of individual loci did not change significantly over the cell cycle (Movie S2), except for loci near *ter*. For a majority of the cell cycle, *ter*-adjacent loci were positioned at the cell center, but at a certain stage, they moved outward and then returned to the center. To highlight this radial repositioning, we combined the mean radial coordinates of 8 selected loci more than 2.06 Mbp away from *oriC*, and plotted them as a function of cell area (Fig. 4c). Notably, for the *ter*-adjacent loci, the radial movement occurred around the same cell cycle stage as the longitudinal movement from the new pole to the middle of the cell.

### A data-driven dynamic polymer model of a replicating chromosome

After acquiring dynamic 3D localization data for 68 loci, we used this data to construct a dynamic polymer model of the *E. coli* chromosome. Each copy of the *E. coli* chromosome was coarse-grained and modeled as a ring of 1,000 beads on a string, with harmonic bonds between neighboring beads and angular harmonic bonds to model the stiffness of the DNA molecule. The chromosome was confined within a cylindrical boundary representing the *E. coli* cell membrane. To inform the polymer model with our imaging data, each bead that corresponds to a labeled locus was tethered by a harmonic force to the mean long axis and radial coordinates obtained from the imaging data. In addition, since we saw a clear global dynamic on the long axis (Fig. 3), we tethered unlabeled beads with a weaker harmonic force to a long axis coordinate based on interpolation between the two neighboring labeled loci (Materials and Methods). We also introduced a weak radial harmonic potential to all beads that correspond to unlabeled loci centered at one-fifth of the cell’s radius to prevent untethered loci from being pushed to the cell’s membrane. The model was implemented based on polychrom (Imakaev, Goloborodko, and Brandão 2019), an existing modeling framework for chromosome polymer models that is based on openMM for molecular dynamics simulations (Eastman et al. 2017).

To model the chromosome during replication and cell growth, we binned the demultiplexed imaging data by cell area using 30 bins, starting from the average birth area and ending with the average division area. In our experiment’s growth conditions, replication is already ongoing at cell birth, i.e., loci close to the *oriC* have two copies at birth and four copies before division. To model the replication process, we initialized the polymer model simulation with four copies of the chromosome where each bead that has not yet been incorporated into the chromosome polymer via replication got a zero mass and no volume. We first initialized the simulation according to the mean loci coordinates at birth, then we ran the simulation for a set number of time steps for each area bin (see Materials and Methods for time scale calibrations). For each new bin, we updated the length of the cylindrical confinement to represent cell growth, updated the input imaging-based spatial coordinates to which the beads are tethered, and assigned mass and volume to newly replicated beads. The progress of the replication fork as a function of cell area was determined based on previously published work, where replication initiates at 2.05 μm^2^ and progresses with a constant rate (Fig. S3) (Gras, Fange, and Elf 2024).

By running an ensemble of 100 independent instances of the model, we reproduced loci location distribution dynamics (Fig. 5). This also implies that the polymer model successfully reproduces chromosome segregation (Movie S3). To check the predictive power of this model, we left out the experimental constraints of 10% of the input loci and compared the simulated long and radial axis distribution dynamics for the left out loci between the simulated and measured localization data using 2D Pearson correlation between the dynamic distributions. We performed this analysis for all 68 loci in 10 different ensembles, where in each ensemble a different set of loci was left out. We observed a mean Pearson correlation coefficient of 0.833 between simulated and measured long axis coordinate dynamics, and 0.62 for a comparison between the radial coordinate dynamics (Fig. 5d). Using randomized localization data that preserves chromosome segregation as input for the model, the resulting distribution of Pearson correlation coefficients for the long axis distribution is significantly lower than the correlation coefficients between the measured data and simulated data (Fig. S4, Wilcoxon rank-sum test, *p* < 10^−17^). However, for the radial coordinate distribution we observe no significant difference between correlation coefficients with the randomized input model, which suggest that loci-specific radial localization data does not contribute to the predictive power of the model (Fig. S4, Wilcoxon rank-sum test, *p* = 0.86).

**Fig. 5:**
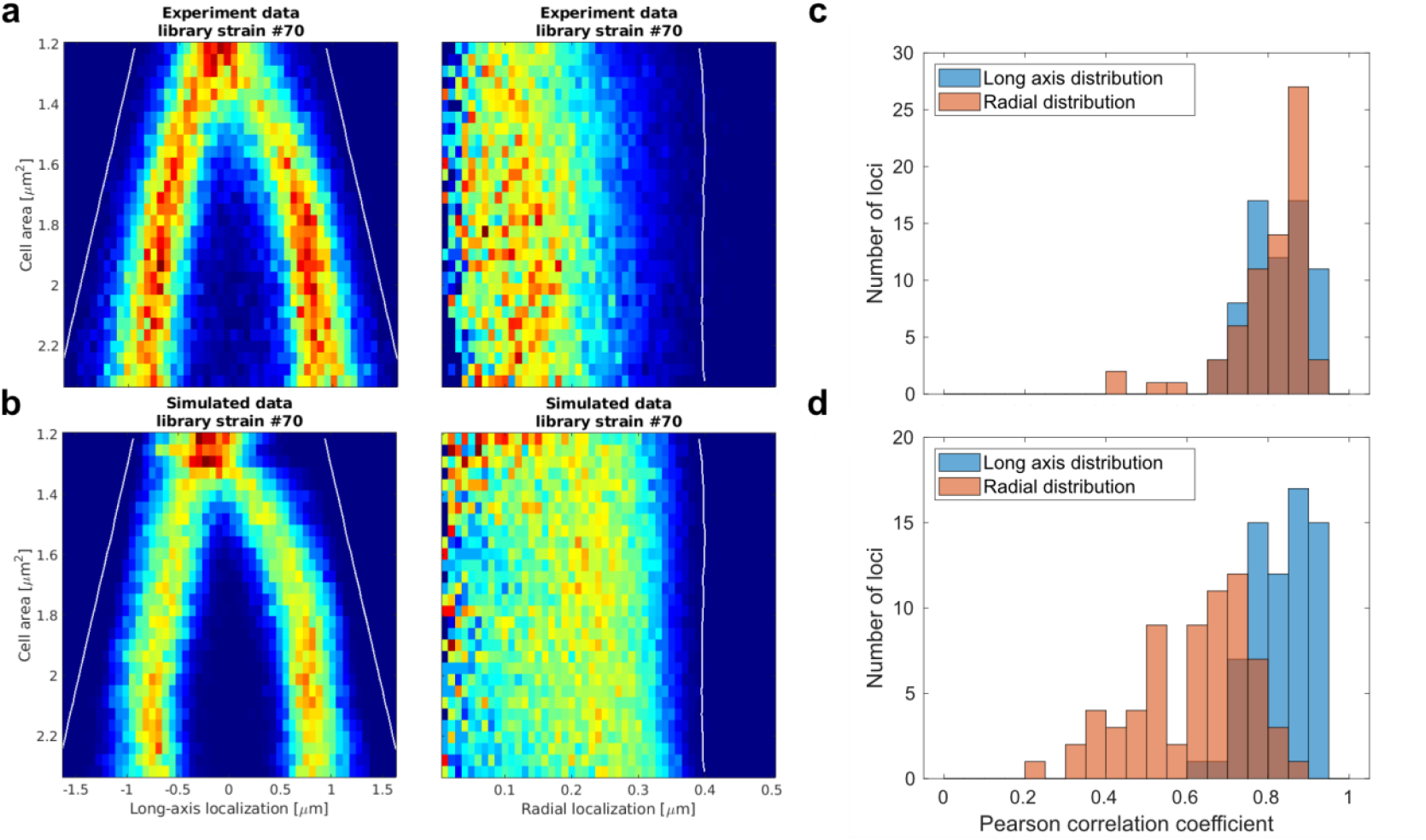
The polymer model of the chromosome structure captures experimental locus locations. **a**, Experimental and **b**, simulated long axis and radial axis location distributions from the polymer model for a selected chromosomal locus. Pearson correlation coefficients between experimental and simulated location distributions were 0.941 for the long axis and 0.686 for the radial axis. Histograms of Pearson correlation coefficients between experimental and simulated long and radial axes coordinate distributions when **c**, all loci location distributions are used as input to the polymer model of the chromosome and **d**, when 10% of the input loci are left out from the model, all input loci were left out by running different ensembles in each of them a different set of 10% was left out. The correlation coefficient is calculated for each locus from the ensemble in which it was left out.

### Modeling of the chromosome structure reveals dynamic macrodomain formation

Using our polymer model, we calculated pairwise distances between loci and visualized them as a pairwise distance map (Fig. 6). The bulk distance map, with 3D distances averaged over all cell areas, indicated two Mbp-sized sequence regions with many short distance interactions around *oriC* and *ter* (Fig. 6a) which suggests a macrodomain level organization of the chromosome. However, we note that while the interaction region around *ter* coincides with the known terminus macrodomain, the region around *oriC* is wider than the known *oriC* macrodomain (Valens et al. 2004; Espeli, Mercier, and Boccard 2008).

**Fig. 6:**
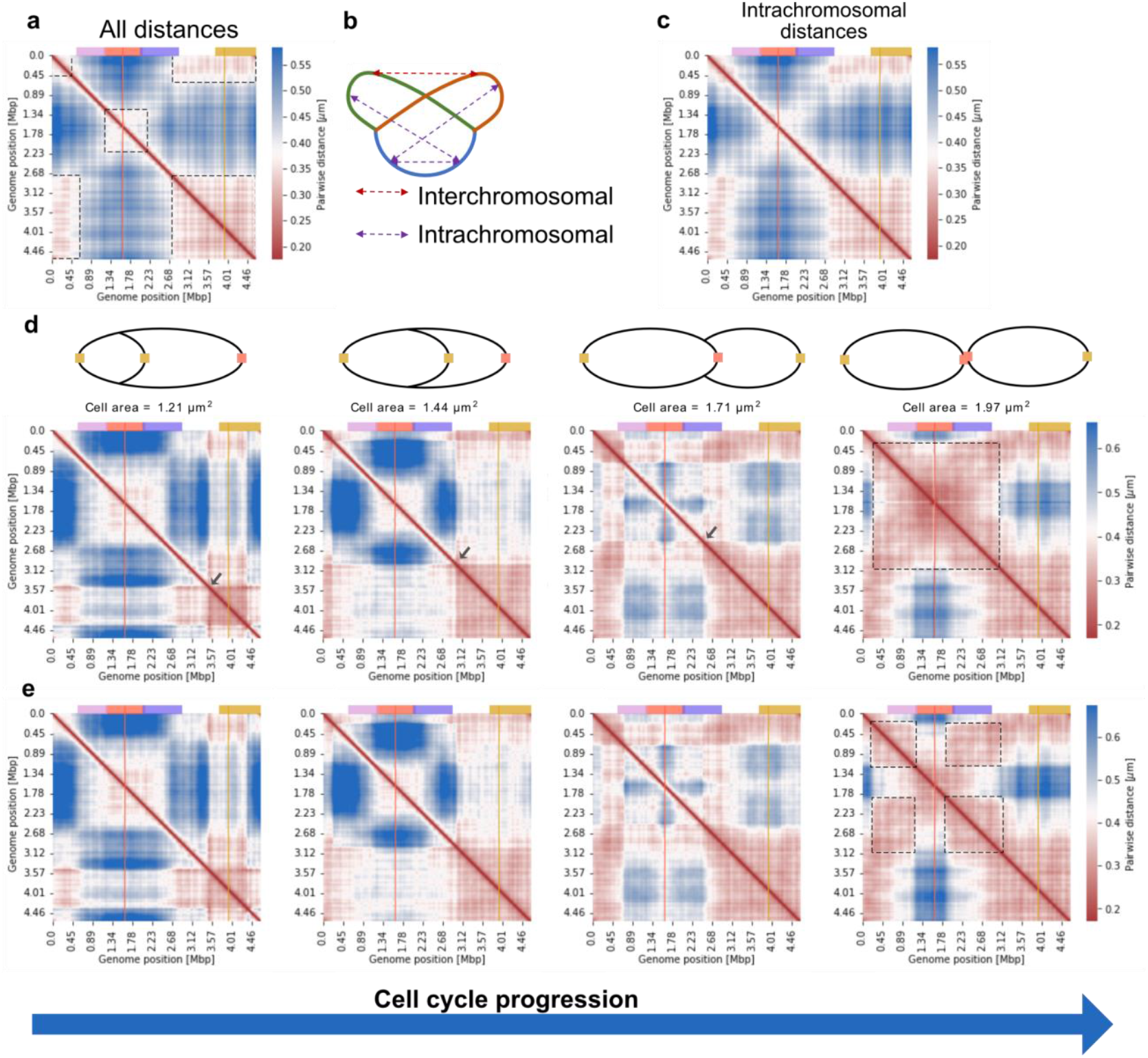
Dynamic distance maps generated with the polymer model of the chromosome. Distance maps averaged over the cell cycle with **a**, all pairwise loci distances and **c**, intra-chromosomal pairwise loci distances. The dashed boxes in a, highlight domains around *oriC* and *ter*. **b**, Cartoon illustrating interchromosomal and intrachromosomal distances. **d**, Distance maps over the cell cycle. As in c, but as a function of cell area (bottom). The grey arrow points to the progression of the replication forks. The dashed box in the rightmost panel highlights a domain that covers the left, right and *ter* macrodomains. (Top) The cartoons illustrate the chromosome in cells with the indicated cell areas. The yellow and red squares show *oriC* and the locus on the opposite side of the chromosome. **e**, Intrachromosomal distance map over the cell cycle. As in d, but for intrachromosomal distances. Each distance map corresponds to a specific cell area bin. The dashed boxes highlight domains that coincide with the macrodomains, and domains that are based on interactions between the left and right chromosome arms. The *oriC*, left, right and *ter* macrodomains are indicated in yellow, blue, pink and red respectively, based on (Espeli, Mercier, and Boccard 2008). The yellow and red lines correspond to *oriC* and the locus on the opposite side of the chromosome. In all distance maps presented, in the case of multiple possible distances for a pair of loci due to replication, the minimal distance is included.

We also used the model to differentiate between intrachromosomal and interchromosomal distances for loci in the two sister chromosomes. We estimated interchromosomal pairwise distances between replicated loci on different copies of the chromosome as well as intrachromosomal distances between unreplicated loci and replicated loci that reside on the same chromosomal molecular ring (Fig. 6b). The interchromosomal distance map (Fig. S5) was in line with the average localizations in (Fig. 2); following replication, the locus copies segregate and move away from each other, except for *ter*-adjacent loci, which remain in the middle. The intrachromosomal distance map (Fig. 6c) showed similar patterns to the all-distance map (Fig. 6a), but with the presence of the *ter* and *oriC* macrodomains being clearer.

The dynamic chromosome structure model made it possible to investigate pairwise loci distances over the cell cycle by binning the distance maps by cell area. Fig. 6c and 6d show all pairwise distances and pairwise intrachromosomal distances respectively at selected cell areas. In both cases, the region around the *oriC* macrodomain exhibited relatively short distances early in the cell cycle. As the cell cycle progressed, the region covered a larger part of the chromosome. The dynamic distance maps showed a momentary decrease in the distances that progressed from *oriC* to *ter*, which we interpreted as a result of replication progression (Fig. 6c, grey arrow). Late into the cell cycle, the all-distance map showed the formation of a macrodomain centered around *ter* that extended towards the macrodomains of the left and right chromosome arms (Fig. 6d, magenta and purple bars). In contrast, the intrachromosomal distance map (Fig. 6e) showed the presence of three smaller macrodomains late in the cell cycle, which overlapped with the macrodomains of the left arm, right arm, and *ter* reported by (Espeli, Mercier, and Boccard 2008). This distance map also showed the presence of a domain found away from the main diagonal based on interactions between the left and right chromosome arms, which is absent in the averaged map (Fig. 6b) and in experimental Hi-C maps (Lioy et al. 2018; Cockram et al. 2021). We also assembled dynamic distance maps with all, intra-, and interchromosomal distances over 30 cell area bins into Movies S4, S5 and S6 to visualize the distances over the cell cycle.

Notably, the dynamic intrachromosomal distance maps showed that the macrodomains form at certain stages of the cell cycle and are not present throughout. The fact that the left and right arm macrodomains appear in our model when considering only intrachromosomal interactions suggests that macrodomain formation is an intrachromosomal property and not a result of interactions between two sister chromosomes. Furthermore, the macrodomains appear to form over the cell cycle as replication progresses, indicating that they may be a result of replication and segregation of the *E. coli* chromosome. To check if chromosome segregation is sufficient for macrodomain formation, we ran the model with randomized input coordinates that maintain chromosome segregation as the centers of the tethering harmonic forces. This was done by randomly assigning an input coordinate for each locus at each cell size bin and to maintain chromosome segregation a coordinate was randomly chosen within the same part of the cell as the measured coordinate from the experimental data (old pole, middle, new pole). However, the resulting dynamic intrachromosomal distance map lacked the left and right arm macrodomains observed in the rightmost panel of Fig. 6d (Fig. S6). This suggests that there are further cellular processes involved in macrodomain formation which are reflected in chromosomal loci dynamics.

## Discussion

We present a dynamic 3D model of the *E. coli* chromosome structure with ∼54 kb resolution. This computational polymer model is based on experimental microscopy data of a library with 68 *E. coli* strains, each with a different locus fluorescently labeled.Combining 3D phenotyping of a strain library with demultiplexing through *in situ* genotyping in a microfluidic device allows us to examine the chromosome over the cell cycle, revealing dynamics that would be obscured using other available methods. The assembled localization data showed global rearrangements of chromosomal loci along the long axis (Fig. 3) that depended on DNA replication, segregation (Gras, Fange, and Elf 2024), and cell division (Espéli et al. 2012). The radial localization data showed two regions located away from the radial center on the left and right chromosomal arms at equal distances from the *oriC* (Fig. 4a and 4b). While most of the loci did not exhibit major radial cell cycle-dependent rearrangements (Movie S2), we observed the average radial positioning of the *ter*-region near the cell center during cell birth, followed by its movement outward and back to the center. At the same time, this region moved from the new pole to the middle of the cell along the long axis (Fig. 4c). The polymer model of the chromosome allowed for the investigation of chromosome dynamics using distance maps (Fig. 6), and also for distinguishing between intrachromosomal and interchromosomal chromosomal distances. The dynamic intrachromosomal distance maps (Fig. 6d) revealed the formation of macrodomains during the cell cycle, which has not been observed previously.

Loci in the *ter* region are positioned at the radial cell center when they are longitudinally located at the new cell pole, or the middle of the cell (Fig. 4c). These 3D localization patterns could be a consequence of the interactions between *ter* and a number of cell division proteins, including MatP (Espéli et al. 2012). Localizations of fluorescently labeled MatP along the cell’s long axis show patterns very similar to *ter* (Sadhir and Murray 2023), starting at the new cell pole and then relocating to the middle. It has been proposed that a MatP-divisome complex can tether *ter* to the cell membrane (Espeli, Mercier, and Boccard 2008). However, while this model may be in line with the radial *ter* localization at the cell center, it does not explain the outward movement. It may be that when *ter* is at the pole or the longitudinal middle, it is tethered by MatP (Espéli et al. 2012), but when *ter* moves from the pole to the middle it becomes untethered and springs outward, away from the cell center.

In the dynamic distance maps, late in the cell cycle (Fig. 6b and 6d), there are domains present away from the diagonal which indicate interactions between the left and right chromosome arms. These domains are less clear in averaged distance maps (Fig. 6a and 6c), and they are also absent from Hi-C contact maps. It has been argued in (Lioy et al. 2018) that the absence of an antidiagonal signal in Hi-C contact maps for the *E. coli* chromosome shows a lack of interaction between the left and right arms, which is present in the contact maps of *Caulobacter cresentus* (Le et al. 2013) and *Bacillus subtilis* (Marbouty et al. 2015). The domain structure shown in our dynamic distance maps late in the cell cycle suggests that the interactions between the left and right arms only occur at certain stages of the cell cycle. Alternatively, the interactions observed in our model could be an artifact of our method, as we do not measure the locations of two different loci in the same cell. We clearly observe that loci on opposing chromosomal arms, with similar distance from *oriC*, share the same long axis and radial mean coordinates (Fig. 2). However, each arm can still have different angular coordinates and occupy opposite sides of the cell, providing the same distributions although there is no interaction between the arms.

Our radial location distributions show that some loci are located away from the radial cell center for most of the cell cycle. This result may seem to present a discrepancy with our previous study, where we report that chromosomal loci move towards a confined replisome region to be replicated (Gras, Fange, and Elf 2024). We have not shown 3D localization distributions of the replisome, but inferring from its relatively narrow localization distribution along the cell’s short axis, we expect the replisome region to be radially localized around the cell center. Thus, we expect loci to sequentially move to the cell center along the radial axis, but we do not observe this in Movie S2. The radial movement of a locus to the replisome at the cell center could be rapid and invisible to us due to the time resolution of our data. Furthermore, we bin localization distributions over cell area, but perhaps the radial locus movement to the replisome is not coordinated with the size of the cells, causing these events to disappear in the population distributions.

While our model successfully predicts longitudinal loci coordinates, and performs well in predicting radial loci coordinates, it doesn’t reproduce known chromosome interaction domains (Lioy et al. 2018) at the 10-100 kb scale, possibly due to the low genomic resolution in this strain library. Solving this by introducing more locus labels would not be possible, as introducing the *malO* array at every locus would disrupt essential genes or affect important regulatory sequences, which could produce mutant phenotypes or kill the cells. To develop a future model of the *E. coli* chromosome structure, we aim to extend the approach presented here to two locus labels, one based on the *malO*/MalI-SYFP2 system and a second locus label based on a different FROS, both in the same cell. We will estimate >5000 3D distances between the two locus labels introduced at a number of different loci in a strain library, with chromosomally expressed barcodes for each locus label. The interlocus distances measured over the cell cycle will inform a new computational model of the chromosome structure. The advantage of measuring distances between two positions instead of absolute positions is that the pairwise distances are independent of the orientation of the bacteria.

With an accurate chromosome structure model, we will be able to study the factors that govern the dynamic organization of the chromosome, e.g. in different growth conditions or with mutant strain backgrounds. It would provide a method to investigate how the DNA sequence information contributes to the structure of the chromosome. We could also study the impact of the dynamic chromosome structure on various functions in *E. coli*. Developing this method will ultimately bring us closer to understanding the functional importance of the bacterial chromosome structure.

## Materials and Methods

### Library construction

To construct the strain library we first constructed plasmid EL3395 containing a chromosomal label cassette which includes the *malO* operator sequence array, a selection marker (*kanR*), and the 3’ backbone for barcode expression. We synthesized an array of 12 repeats of the *malO* operator separated by a random sequence of 15 nucleotides (GenScript), and cloned this array together with a kanamycin resistance gene taken from plasmid pKD4 and the 3’ end of the barcode expression gene (as described in (Soares et al. 2023)) into a pR6K vector using NEBuilder HiFi DNA assembly Master Mix (NEB #E2621L) (see list of oligos in Table S2). The plasmid was based on a pR6K vector which requires a *pir+* cloning strain for plasmid replication, to prevent unintentional carryover of the plasmid during the final strain construction.

Each library strain was constructed individually using lambda red recombination (Zhang et al. 1998). For all library strains, the label cassette from EL3395 was inserted into the chromosome of a background strain EL3420 that expresses the MalI transcription factor fused to SYFP2 (*E. coli* MG1655 *rph*+ *ΔmalI::frt intC::P59-malI-SYFP2-frt gtrA::P58-mCherry2-parB-SpR*). For each strain, the cassette was PCR amplified by a unique primer pair, where each of them included 40 nucleotide homology sequences to the desired insertion site. The 5’ primer also included a unique barcode sequence and the 5’ backbone of the barcode expression downstream of the chromosomal homology sequence. Barcode sequences were picked from a set of previously verified barcodes as described in (Soares et al. 2023). The amplified cassette was transformed into EL3420 cells using lambda red recombination and selected on LB agar plates with kanamycin. In choosing library insertion sites we aimed for even coverage of the *E. coli* chromosome while choosing insertion sites that are predicted to have minimal effect on cellular fitness. We favored first pseudogenes, second intergenic regions between two genes in opposing orientations such that the insertion site will be at the 3’ end of both genes (lower probability for regulatory elements to be found there), and third if none of the above was found in a desired region we chose a coding gene. All insertion sites were corroborated using high throughput TnSeq (Price et al. 2016) and CRISPR-dCas9 (Calvo-Villamañán et al. 2020) screens to check that insertions in these sites are predicted to cause no growth defects (see Table S3 for a list of all insertion sites, primers, and barcode sequences).

To avoid off-target mutations due to lambda red recombination we transferred the inserted label cassette back into EL3420 cells using P1vir phage transduction (Thomason, Costantino, and Court 2007). At this step, strains were pooled together and transduction was done in sub-pools of 10-11 strains each. To avoid biases resulting from different target size due to replication, strains were pooled together according to the distance of the label insertion site from *oriC* (Masters 1977). For each sub-pool we pooled stationary phase overnight cultures grown in LB media of the different strains at equal volume, prepared a phage lysate from that pool, and transduced it into EL3420 cells. After transduction the acceptor cells were plated on a LB agar plate containing kanamycin and 10% sodium citrate. To ensure the removal of the P1 phage we collected all the colonies from the plates and replated on LB agar plates at 3 different concentrations (10^−6^, 10^−7^, 10^−8^ cells/ml), and the plate that yielded an order of 100 colonies was selected and all the colonies were collected. After transduction in sub-pools, all resulting sub-pools were grown overnight in LB to stationary phase and pooled together at volumes proportional to the number of strains in each sub-pool to create the final 83 strains library.

### Experiment conditions

#### Phenotyping

The phenotyping experiments were performed in M9 minimal medium supplemented with 51 μg/ml Pluronic F-108 (Sigma-Aldrich 542342) 0.4% succinate and 1× RPMI 1640 amino acid solution (Sigma) at 30 °C. The strain library was inoculated one day before each experiment in culture tubes with growth medium from frozen stock cultures stored at −80 °C. Cells were grown overnight at 30 °C in a shaking incubator (200 rpm). On the day of the experiment, the cell culture was diluted 1:100 in growth medium and grown for 3 to 4 h before being loaded in the microfluidic chip (Baltekin et al. 2017).

A mother machine-type PDMS chip with open-ended channels was used for all microfluidic experiments. The width and height of these channels was 1000 nm. Medium flow and loading of cells into the chip was performed by supplying pressure to the chip ports with a microfluidic flow controller built in-house (AnduinFlow). The imaging of the chip was performed in a H201-ENCLOSURE hood that enclosed the microscope stage, with the hood being connected to a H201-T-UNIT-BL temperature controller (OKOlab).

#### Genotyping

The genotyping protocol was performed in the same microfluidic chip following phenotyping. Cells were permeabilized by exchanging port 2.0 (Baltekin et al. 2017) consisting of M9 minimal growth medium with 70% EtOH and flown into the sample for 30 min at a pressure 200 mBar. A counter pressure at 40 mBar was applied at the cell loading ports used during the phenotyping experiment (ports 2.1 and 2.2, as described in (Baltekin et al. 2017)). Subsequently, all connected ports were then carefully replaced with 1x PBS buffer with 0.1% Tween (PBST). The cells were rehydrated with PBST from port 2.0 at 200 mBar, overnight at room temperature.

Next, cells were partly degraded with lysozyme (250 mg/mL) using a syringe pump at a flow rate of 1 μl/min and with a counter pressure 40 mBar at ports 2.0, 2.1, and 2.2 respectively. The lysozyme activity was then blocked by flowing in bovine serum albumin with Tween for 10 mins at 1 μl/min flow-rate. Hereafter we exchanged the reaction mixtures by plugging and unplugging at port 2.0, maintained 30 °C and applied a counter pressure of 40 mBar, until the last round of genotype imaging.

Details of the reaction mixes used for *in situ* genotyping can be found in Table S4. Barcode expression was performed by applying the transcription mix at a 0.5 μl/min flow-rate for 2 hours. Then, padlock probes (PLP) were hybridized to the expressed barcodes by flowing in the PLP hybridization mix at a 1 μl/min flow-rate for 30 min. This was followed by ligation of the PLPs with the SplintR ligation mix at a 1 μl/min flow-rate for 30 min. After the ligation, the resulting circularized product was amplified with rolling circle amplification (RCA), using the RCA mix flown in at a 0.5 μl/min flow-rate for 2 hours. Finally, the RCA products were detected with fluorescently labeled detection oligonucleotides in the labeling mix, and flown in at 1 μl/min flow-rate for 30 min.

Barcode detection: In each sequential hybridization round, a mix of adaptor “L-probes” were hybridized to the RCA products, followed by parallel hybridization with 4 differently coloured fluorescent detection probes. For each round of sequential FISH, the L-probes, which were pre-hybridised to detection oligos, were flown onto the chip at 1 μl/min for 30 min followed by imaging for genotyping. Probes were removed from the RCA product by flowing the probe stripping mixture at 1 μl/min for 30 min. Probe hybridization, imaging, and stripping steps were repeated an additional 6 times with different known variations of L-probe detection oligo pools according to the genotyping code.

### Microscopy

#### Phenotyping

Imaging of the phenotyping experiments was performed with a Ti-E (Nikon) microscope equipped with 100X immersion oil objective (Nikon, NA 1.45, CFI Plan Apochromat Lambda D MRD71970CFI) for phase contrast and widefield epi-fluorescence microscopy. Phase contrast and fluorescence images were acquired of cells growing in a microfluidic chip at 30 °C in M9 minimal medium supplemented with 0.4% succinate and 1x RPMI amino acids solution (Sigma). The imaging was performed over 10 h, unless otherwise noted. Phase contrast images were acquired once per position on the microfluidic device with a 2 min^-1^ imaging frequency. Fluorescence images were acquired once per position with a 4 min^-1^ imaging frequency. The imaging was performed using in-house plugins in Micro-Manager.

For fluorescence image acquisition, the sample was irradiated with a 515 nm laser (Fandango 150, Cobolt) for SYFP2 excitation. The laser was set to a 25 ms exposure time and 30 W/cm^2^ power density at the image plane. The laser light was shuttered using an AOTFnC together with MPDS (AA Opto-Electronic) and reflected onto the sample using a FF444/521/608-Di01 (Semrock) triple-band dichroic mirror. Fluorescence images were acquired using a Kinetix sCMOS camera (Teledyne Photometrics). Imaging of the fluorescence channel was performed by a function generator (Tektronix) that triggered the laser based on the camera acquisition. Emitted fluorescence was transmitted through an astigmatic lens, followed by a BrightLine FF580-FDi02-T3 (Semrock) dichroic beamsplitter to separate the fluorescence for different channels. The fluorescence was then filtered through a BrightLine FF01-505/119-25 (Semrock) filter and focused on the sCMOS camera.

Phase contrast images were acquired with a 25 ms exposure time using a DMK 38UX304 camera (The Imaging Source). The light source used for phase contrast was a 480 nm LED and a TLED+ (Sutter Instruments). The transmitted light was passed through the same FF444/521/608-Di01 (Semrock) triple-band dichroic mirror as the fluorescence and reflected onto the camera with a Di02-R514 (Semrock) dichroic mirror. Phase contrast images were acquired by directing the light transmitted through the microfluidic sample onto a different camera than the one used for fluorescence image acquisition, thus avoiding transmission through the cylindrical lens.

#### Genotyping

Imaging of the genotyping experiments was performed using a Ti2-E (Nikon) microscope for both phase contrast and widefield epifluorescence. The microscope was equipped with a 100x CFI Plan Apo Lambda DM (Nikon) objective and images were acquired using a Sona camera (Andor). Phase contrast images were acquired with 80 ms exposure time and fluorescence images were acquired with 300 ms exposure time. Temperature was maintained using a temperature controller and hood (OKOlab), which enclosed the microscope stage. The light source used for fluorophore excitation was a Spectra III (Lumencore). The following light source and dichroic mirror combinations were used for each detection oligonucleotide fluorophore: cyan channel with EX450-490 (Nikon) for excitation and FF01-524/24 (Semrock) for emission of Alexa488, green channel with FF01-543/22 (Semrock) for excitation and FF01-586/20 (Semrock) for emission of Cy3, red channel with FF660-Di02 (Semrock) for excitation and 692/40 (Semrock) for emission of Cy5, and NIR channel with T760lpxr (Chroma) for excitation and ET811/80 (Chroma) for emission of Cy7.

### Image analysis

#### Phenotyping

The image analysis was performed using an automated image analysis pipeline that was primarily written in MATLAB R2022a (Mathworks) and previously described in (Camsund et al. 2020). Cell segmentation of phase contrast images was performed using the Omnipose method (Kandavalli et al. 2022; Cutler et al. 2022). Cell tracking was performed using the Baxter algorithm (Magnusson et al. 2015). Fluorescent SYFP2 foci were localized in 3D using a field-dependent DeepLoc algorithm (Karempudi et al. 2023). To transform the localized focus coordinates between the cameras used to acquire fluorescence and phase contrast images, landmark-based registration was performed using 500 nm beads (TetraSpeck, Thermo Fisher) that were visible on both cameras.

The transformed focus coordinates were converted into internal long and short axis coordinates using a cell backbone fit (Wallden et al. 2016). The long axis coordinates were binned along the cell’s length, with zero corresponding to the middle of the cell, resulting in a histogram over all long axis coordinates. The histograms were then normalized by the number of foci per cell. To generate the long axis coordinate distributions (Fig. 3 and S2), the histograms were binned as a function of the cell’s area.

The short and depth axis coordinates were used to generate the radial axis distributions. The depth axis was relative to the average z-coordinate over the image plane for each field of view. The estimated radial coordinates were binned along different radial positions of the cell, resulting in radial histograms. These histograms were then binned as a function of the cell’s area. To avoid introducing a bias by sorting coordinates into bins with different spatial areas, we normalized the radial histograms by the area of each radial bin. We then binned the normalized radial histograms as a function of cell area to generate the radial location distributions (Fig. 3 and S2).

#### Genotyping

Phase contrast images from each round of genotyping were aligned using the dot barcodes imprinted next to empty growth channels. Growth channel segmentation was performed using the deep learning model described in (Kandavalli et al. 2022). Empty channels were detected by identifying channels that include no cells’ lysate. As empty channels have a smooth image compared to a rugged pattern where cells’ lysate exists, empty channels were identified by thresholding the magnitude of the highest bins in a Fourier transform of the cropped channels images. Fluorescent blobs were detected using a signal threshold that was five standard deviations higher than the mean of the Gaussian fitted to the histogram of pixel intensities from all of the growth channels in each position in the same fluorescence channel. The resulting fluorescence masks were compared between rounds and the final mask consisted of the pixels that appear at least in half of the rounds. For the growth channel code assignment, the fluorescent signal from a channel was considered only if (i) it passed the intensity threshold, (ii) the number of pixels passing the intensity threshold passed an area threshold, and (iii) the signal was within the accumulated fluorescence mask. If after applying the thresholds and the mask there was a signal in a single channel, it was designated as the decoded channel. Otherwise if there was a signal in multiple channels, a channel was designated as the decoded channel only if its total intensity was at least twice higher than all other channels. We applied error correction to growth channel codes fixing at most 1 round, by assigning the decoded sequence to one of the code words if it was identical to this code word, or one error away from it (i.e. Hamming distance equals 1) (Chen et al. 2015).

### Genotyping code design

A pool of 97 L-probes (The L-probes pool was designed for a bigger strain library) was mixed for each hybridization-round according to the code allowing for error correction (see Table S5 for a list of the L-probes used). Each strain was assigned a randomly chosen 7-symbol code word (with an alphabet of 4 symbols), while ensuring that each code word has a Hamming distance of at least 3 symbols from all other 96 code words. Each symbol in the code is represented by a differently colored detection oligo. Thus, for each hybridization-round we mixed a collection of 97 L-probes each connecting the padlock probes readout barcode and the correct detection oligo for that round.

To support a phenotypic outlier detection step (see below) we wished that a strain will have a lower probability to be misdetected as another strain with a similar phenotype. We predicted that two strains with a similar distance from *oriC* would present a similar phenotype (Fig. 2).

Therefore, each strain was assigned with a 7-symbol code word with a Hamming distance of at least 4 symbols away from all other strains with a label insertion site at a distance from *oriC* which is within a range of 500,000 base pairs from that strain’s distance from *oriC*.

Meaning for each strain *s* a distance of 4 symbols was assured against all strains *x* that fulfill the following:

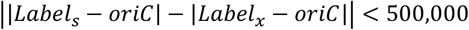

### Phenotypic outlier detection

The output of a genotyping image analysis is a list of strain identities assigned to growth channels on the chip. Using pooled data from 4 independent imaging experiments we collected all growth channels assigned to each strain and first produced the pooled long axis distribution as a function of cell area per strain as in Fig. 2. If a strain was observed in less than 10 growth channels we discard it. Next, to discard growth channels that were misdetected during genotyping analysis, we assume that the probability of misdetection is low enough such that the pooled distribution across all growth channels assigned to a strain faithfully represents the strain’s label long-axis distribution phenotype. We further assume that the long axis distribution is similar for strains with a label at a similar distance to *oriC*. As explained above, the genotyping code was designed such that misdetection of a strain with a similar phenotype is less probable.

To detect phenotypic outlier growth channels we did the following per growth channel *i*.

1. Produce the long axis two dimensional distribution for a single growth channel *i* assigned to strain *s*_*genotyped*_.
2. Compute *r*_*i*_(*s*), the Pearson correlation coefficient of the single channel distribution against all strains’ pooled distribution.
3. Find the strain that maximizes the correlation vector *s*_*max*_= *argmax* (*r*_*i*_(*s*)).
4. Discard channel *i* if one of the following conditions is true:

a. The difference in the strain’s label distance from *oriC* between *s*_*max*_ and the genotyped strain *s*_*genotyped*_ is more than 0.02 ⋅ {*genome size*}.

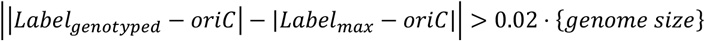
b. The correlation coefficient of the genotyped strain is more than 0.05 away from the maximal correlation.

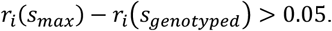

### Amplicon sequencing

To measure the relative frequency of each strain in the library, we grew a culture of the library pool in the same growth conditions as for the phenotyping experiment (M9 minimal media + 20% succinate + amino acids, at 30 °C). 3 samples of 0.5 ml were collected for 3 technical repeats. Chromosomal DNA was extracted using Monarch genomic DNA purification kit (NEB #T3010S). The barcode region was amplified with primers prDS292-prDS293 which include Illumina sequencing adaptors (Phusion HiFi mastermix standard protocol, Thermo Fisher #F531S). The PCR product was further amplified with TruSeq UDI indexed primers 4-6, to uniquely index each sample (Phusion HiFi master mix, 62 °C annealing temperature, 15 cycles). The 3 samples were pooled and sequenced in an Illumina iSeq 100 according to the manufacturer’s recommended protocol. The sequencing reads were matched to barcode sequences using bowtie2 (Langmead and Salzberg 2012) and the relative frequency of each barcode was counted in each repeat. Pairwise Pearson correlations between the 3 repeats were higher than 0.99, so we took the mean frequency over the 3 repeats.

### Modeling and simulations

#### Polymer simulation and parameters

The chromosome is modeled as a beads-on-a-string polymer model using polychrom (Imakaev, Goloborodko, and Brandão 2019), a framework for chromosome polymer simulations that is based on OpenMM for molecular dynamics simulations (Eastman et al. 2017). Each monomer in the simulation has a diameter of 1 nm and a mass of 100 Daltons in simulation space, all other units are scaled accordingly. Each monomer is associated with an excluded volume potential using polychrom’s default potential with a truncation parameter that equals √1.5, which allows chains to cross each other. Each neighboring bead is connected through a harmonic bond using polychrom’s default parameters, and the polymer’s stiffness is simulated using an angular harmonic bond with a spring coefficient K=2. The polymer resides within a cylindrical confinement implemented using polychrom’s function with K=1, the dimensions such that its radius is half of its length, and monomers that form a single copy of the chromosome occupy 10% of the cylinder’s volume at the start of the simulation (before cell growth).

A single copy of the chromosome comprises 1000 monomers. Since in our experiments, there are 4 copies of the chromosome at least partially replicated at cell division, we initialized the simulation with all possible copies, meaning 4000 monomers. Only monomers that are already replicated at cell birth are assigned with a mass and excluded volume potential.

#### Extracting coordinates from distribution plots

To inform the simulation with data-driven coordinates per locus, we needed to extract representative long-axis and radial coordinates from the distributions for each cell size bin. We binned the cell sizes to 30 bins, from the mean cell size at birth to the mean cell size at division. For the radial coordinate distribution, for each cell size bin, we extracted a histogram of radial locations over equally sized radial bins and normalized it by the relative area of each radial bin. For the radial distribution, we took the mode of the distribution and its standard deviation at each cell size bin, to represent the radial coordinate. For the radial coordinate, we assumed that all copies of the same locus have the same radial localization (Fig. S7).

For the long-axis coordinate we first needed to decompose the distribution to the different chromosome copies. We assessed the number of expected chromosome copies based on the replication fork progression as a function of the locus’ distance from *oriC* and the cell size. This function was modeled based on previously published data where the replisome was tracked for 12 of this libraries strains and the cell size in which replication occurs was identified ?(Gras et al. 2024) (Fig. S3). Coordinates were extracted depending on the number of chromosomal copies the following way:

1. Single copy - The distribution was fitted to a single gaussian model and the mean and standard deviation of the fitted model were used.
2. Two copies - The distribution was fitted to a two gaussians mixture model, and the mean and standard deviation of each modeled gaussian were assigned to each copy.
3. Four copies - The long axis of the cell was divided to two halves at the cell center, and for each half a two gaussian mixture model was fitted the same as in the case of two copies.
4. Whenever a two gaussian mixture model was fitted, and the second gaussian proportion was lower than ⅓ we assumed that the two copies of this label are co-localized and assigned both copies the same coordinate.

#### Simulating replicating chromosome with imaging-based input

The chromosome polymer simulation we performed is a dynamic simulation of a replicating chromosome in a growing cell.

We first initiliaze the simulation the following way:

1a. The polymer is initialized by localizing monomers that correspond to labeled loci at their measured coordinates at birth.
1b. Monomers that correspond to unreplicated loci are initialized with a zero mass, no excluded volume potential, and are localized next to the already existing copy of the same locus.
1c. Run energy minimization.
1d. Run the simulation to equilibrium (200,000 simulation steps) to avoid any artifacts due to the artificial initial conformation.

After initialization, we run the following steps for each cell size bin in order:

2a. Apply tethering harmonic bonds:

i. If a monomer is associated with a labeled locus from the library: the harmonic bond is tethered to the measured coordinate with a spring coefficient with a long axis and radial component that equal to:

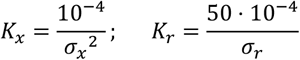
ii. If a monomer is not associated with a labeled locus it is tethered to a long axis coordinate the is determined using linear regression between the two neighboring monomers that are associated with labeled loci, and to *r* = 0.2 ⋅ *R* where R is the cell’s radius. The spring coefficients are given by (*x*1 and *x*2 represent the two neighboring labeled monomers):

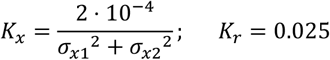

2b. Lengthen the cylinder by the relative change in cell length compared to the previous cell size bin.
2c. Assign mass and excluded volume potential to newly replicated monomers
2d. Move unreplicated monomers to the location of their corresponding replicated monomer.
2e. Run energy minimization
2f. Run simulation for 66,666 simulation steps. The number of simulation steps corresponds to the time spent in each of the 30 cell size bins, assuming a generation time of 1 hour (i.e. 2 minutes per cell size bin). Calibration of simulation time units and real time is described below.

#### Calibration of simulation steps and real-time units

To calibrate the simulation time units with real time we measured the mean square displacement (MSD) of a labeled locus both in the cells and in the simulation. For the experimental measurement, we used the same imaging setup as described above, but without a cylindrical lens. For this purpose, we used a single strain with a label next to *ter* (library strain M35), since it presents a single fluorescent focus through most of the cell cycle. We acquired one phase contrast image with 50 ms exposure time followed by a stack of 10 fluorescence images acquired with 100 ms exposure time at each position on the microfluidic device. To measure locus displacements over different time lags, the fluorescence image stacks were acquired with different unexposed time intervals between each frame, with the exposure time and time interval summing to 100, 150, 200, 500, 1000, 1500, 2000 and 3000 ms time lags. We acquired image stacks for four positions per time lag. The images were analyzed as for the phenotyping experiments, except for the detection and localization of foci, which was done with a wavelet transform (Olivo-Marin 2002) and fitting of a 2D normal distribution function (Lindén et al. 2017). The fluorescent foci were also tracked using u-track (Jaqaman et al. 2008) to assemble trajectories of the loci. The trajectories were post-processed to estimate loci MSD as a function of time lag.

At the simulation space, we measured displacement by running an ensemble of 100 simulations of a single copy of the chromosome confined in a cylinder corresponding to the cell size at birth and taking snapshots at different time lags after first running the simulation to equilibrium. We calculated the MSD values at different time lags for an arbitrarily chosen monomer. To get the calibration factor we plotted both the empirical and simulated MSD curves in log-log scale in which we expected the curve to be linear, and found a scaling factor for the simulated time units that aligns the two curves.

## Data availability

Raw data and analysis output can be accessed in a Figshare repository at https://figshare.com/s/ef5f1fbb7a4a4daf0b6a.

## Code availability

Analysis code for the imaging data used in this study can be accessed at https://figshare.com/s/ef5f1fbb7a4a4daf0b6a. All modeling simulation code and code for some of the figures can be found at https://github.com/DvirSchirman/SingleLabelChrModel.

## Acknowledgements

We thank Irmeli Barkefors for the critical reading of the manuscript; Anton Goloborodko and Ankit Gupta for helpful discussions and advice on the modeling and simulations. Spartak Zikrin developed the analysis pipeline. Praneeth Karempudi trained the 3D localization network. Anna Knöppel was involved in the design of the pooled strain library. Elias Amselem built the optical setup. Madeleine Skeppås and Emma Svanberg Frisinger collected the data and performed the analysis for the simulation time scale calibrations. We thank Hervé Nicoloff for advice on pooled phage transductions. This study was made possible by grants from the ERC (advanced grant no. 885360), the Swedish Research Council (grant nos. 2016-06213 and 2018-03958), the Knut and Alice Wallenberg Foundation (grant nos. 2017.0291, and 2019.0439) and the eSSENCE e-science initiative. D.S. was supported by EMBO postdoctoral fellowship. The computations and data management were enabled by resources provided by the Swedish National Infrastructure for Computing at UPPMAX, partially funded by the Swedish Research Council through grant agreement no. 2018-05973.

## Author contributions

J.E. conceived the study. D.S. and K.G. contributed equally to this work and their order in the author list was decided by lottery. D.S. constructed and designed the pooled strain library, analyzed the genotyping experiments and developed the modeling simulations. K.G. developed the phenotyping, performed all of the phenotyping experiments and analyzed the data. V.K. performed all of the genotyping experiments. J.L. developed the genotyping protocol. D.S., K.G., D.F. and J.E. interpreted results. K.G. and D.S. wrote the paper with input from D.F. and J.E.

## References

Askary, Amjad, Luis Sanchez-Guardado, James M. Linton, Duncan M. Chadly, Mark W. Budde, Long Cai, Carlos Lois, and Michael B. Elowitz. 2020. “In Situ Readout of DNA Barcodes and Single Base Edits Facilitated by in Vitro Transcription.” Nature Biotechnology 38 (1): 66–75.

Badrinarayanan, Anjana, Tung B. K. Le, and Michael T. Laub. 2015. “Bacterial Chromosome Organization and Segregation.” Annual Review of Cell and Developmental Biology 31:171–99.

Baltekin, Özden, Alexis Boucharin, Eva Tano, Dan I. Andersson, and Johan Elf. 2017. “Antibiotic Susceptibility Testing in Less than 30 Min Using Direct Single-Cell Imaging.” Proceedings of the National Academy of Sciences 114 (34): 9170–75.

Bates, David, and Nancy Kleckner. 2005. “Chromosome and Replisome Dynamics in E. Coli: Loss of Sister Cohesion Triggers Global Chromosome Movement and Mediates Chromosome Segregation.” Cell 121 (6): 899–911.

Bignaud, Amaury, Charlotte Cockram, Céline Borde, Justine Groseille, Eric Allemand, Agnès Thierry, Martial Marbouty, Julien Mozziconacci, Olivier Espéli, and Romain Koszul. 2024. “Transcription-Induced Domains Form the Elementary Constraining Building Blocks of Bacterial Chromosomes.” Nature Structural & Molecular Biology 31 (3): 489–97.

Bintu, Bogdan, Leslie J. Mateo, Jun-Han Su, Nicholas A. Sinnott-Armstrong, Mirae Parker, Seon Kinrot, Kei Yamaya, Alistair N. Boettiger, and Xiaowei Zhuang. 2018. “Super-Resolution Chromatin Tracing Reveals Domains and Cooperative Interactions in Single Cells.” Science 362 (6413). 10.1126/science.aau1783.

Calvo-Villamañán, Alicia, Jérome Wong Ng, Rémi Planel, Hervé Ménager, Arthur Chen, Lun Cui, and David Bikard. 2020. “On-Target Activity Predictions Enable Improved CRISPR– dCas9 Screens in Bacteria.” Nucleic Acids Research 48 (11): e64.

Camsund, Daniel, Michael J. Lawson, Jimmy Larsson, Daniel Jones, Spartak Zikrin, David Fange, and Johan Elf. 2020. “Time-Resolved Imaging-Based CRISPRi Screening.” Nature Methods 17 (1): 86–92.

Cass, Julie A., Nathan J. Kuwada, Beth Traxler, and Paul A. Wiggins. 2016. “Escherichia Coli Chromosomal Loci Segregate from Midcell with Universal Dynamics.” Biophysical Journal 110 (12): 2597–2609.

Chen, Kok Hao, Alistair N. Boettiger, Jeffrey R. Moffitt, Siyuan Wang, and Xiaowei Zhuang. 2015. “RNA Imaging. Spatially Resolved, Highly Multiplexed RNA Profiling in Single Cells.” Science 348 (6233): aaa6090.

Cockram, Charlotte, Agnès Thierry, Aurore Gorlas, Roxane Lestini, and Romain Koszul. 2021. “Euryarchaeal Genomes Are Folded into SMC-Dependent Loops and Domains, but Lack Transcription-Mediated Compartmentalization.” Molecular Cell 81 (3): 459–72.e10.

Cutler, Kevin J., Carsen Stringer, Teresa W. Lo, Luca Rappez, Nicholas Stroustrup, S. Brook Peterson, Paul A. Wiggins, and Joseph D. Mougous. 2022. “Omnipose: A High-Precision Morphology-Independent Solution for Bacterial Cell Segmentation.” Nature Methods 19 (11): 1438–48.

Dekker, Job, Karsten Rippe, Martijn Dekker, and Nancy Kleckner. 2002. “Capturing Chromosome Conformation.” Science 295 (5558): 1306–11.

Eastman, Peter, Jason Swails, John D. Chodera, Robert T. McGibbon, Yutong Zhao, Kyle A. Beauchamp, Lee-Ping Wang, et al. 2017. “OpenMM 7: Rapid Development of High Performance Algorithms for Molecular Dynamics.” PLoS Computational Biology 13 (7): e1005659.

Espéli, Olivier, Romain Borne, Pauline Dupaigne, Axel Thiel, Emmanuelle Gigant, Romain Mercier, and Frédéric Boccard. 2012. “A MatP–divisome Interaction Coordinates Chromosome Segregation with Cell Division in E. Coli.” The EMBO Journal 31 (14): 3198– 3211.

Espeli, Olivier, Romain Mercier, and Frédéric Boccard. 2008. “DNA Dynamics Vary according to Macrodomain Topography in the E. Coli Chromosome.” Molecular Microbiology 68 (6): 1418–27.

Fisher, Jay K., Aude Bourniquel, Guillaume Witz, Beth Weiner, Mara Prentiss, and Nancy Kleckner. 2013. “Four-Dimensional Imaging of E. Coli Nucleoid Organization and Dynamics in Living Cells.” Cell 153 (4): 882–95.

Fraser, James, Mathieu Rousseau, Solomon Shenker, Maria A. Ferraiuolo, Yoshihide Hayashizaki, Mathieu Blanchette, and Josée Dostie. 2009. “Chromatin Conformation Signatures of Cellular Differentiation.” Genome Biology 10 (4): R37.

Gras, Konrad, David Fange, and Johan Elf. 2024. “The Escherichia Coli Chromosome Moves to the Replisome.” Nature Communications 15 (1): 6018.

Harju, Janni, Muriel C. F. van Teeseling, and Chase P. Broedersz. 2024. “Loop-Extruders Alter Bacterial Chromosome Topology to Direct Entropic Forces for Segregation.” Nature Communications 15 (1): 4618.

Hua, Kang-Jian, and Bin-Guang Ma. 2019. “EVR: Reconstruction of Bacterial Chromosome 3D Structure Models Using Error-Vector Resultant Algorithm.” BMC Genomics 20 (1): 738.

Imakaev, Maksim, Anton Goloborodko, and Hugo B. Brandão. 2019. Mirnylab/polychrom: v0.1.0. Zenodo. 10.5281/ZENODO.3579473.

Javer, Avelino, Nathan J. Kuwada, Zhicheng Long, Vincenzo G. Benza, Kevin D. Dorfman, Paul A. Wiggins, Pietro Cicuta, and Marco Cosentino Lagomarsino. 2014. “Persistent Super-Diffusive Motion of Escherichia Coli Chromosomal Loci.” Nature Communications 5 (May):3854.

Javer, Avelino, Zhicheng Long, Eileen Nugent, Marco Grisi, Kamin Siriwatwetchakul, Kevin D. Dorfman, Pietro Cicuta, and Marco Cosentino Lagomarsino. 2013. “Short-Time Movement of E. Coli Chromosomal Loci Depends on Coordinate and Subcellular Localization.” Nature Communications 4:3003.

Kandavalli, Vinodh, Praneeth Karempudi, Jimmy Larsson, and Johan Elf. 2022. “Rapid Antibiotic Susceptibility Testing and Species Identification for Mixed Samples.” Nature Communications 13 (1): 6215.

Kao, H. P., and A. S. Verkman. 1994. “Tracking of Single Fluorescent Particles in Three Dimensions: Use of Cylindrical Optics to Encode Particle Position.” Biophysical Journal 67 (3): 1291–1300.

Karempudi, P., K. Gras, E. Amselem, S. Zikrin, and J. Elf. 2023. “Three-Dimensional Localization of Fluorescent Proteins in Living Escherichia Coli.” bioRxiv. https://www.biorxiv.org/content/10.1101/2023.10.20.563292.abstract.

Knöppel, Anna, Oscar Broström, Konrad Gras, Johan Elf, and David Fange. 2023. “Regulatory Elements Coordinating Initiation of Chromosome Replication to the Escherichia Coli Cell Cycle.” Proceedings of the National Academy of Sciences of the United States of America 120 (22): e2213795120.

Langmead, Ben, and Steven L. Salzberg. 2012. “Fast Gapped-Read Alignment with Bowtie 2.” Nature Methods 9 (4): 357–59.

Lau, Ivy F., Sergio R. Filipe, Britta Søballe, Ole-Andreas Økstad, Francois-Xavier Barre, and David J. Sherratt. 2004. “Spatial and Temporal Organization of Replicating Escherichia Coli Chromosomes: Escherichia Coli Chromosome Dynamics.” Molecular Microbiology 49 (3): 731–43.

Lawson, Michael J., Daniel Camsund, Jimmy Larsson, Özden Baltekin, David Fange, and Johan Elf. 2017. “In Situ Genotyping of a Pooled Strain Library after Characterizing Complex Phenotypes.” Molecular Systems Biology 13 (10): 947.

Le, Tung B. K., Maxim V. Imakaev, Leonid A. Mirny, and Michael T. Laub. 2013. “High-Resolution Mapping of the Spatial Organization of a Bacterial Chromosome.” Science 342 (6159): 731–34.

Lioy, Virginia S., Axel Cournac, Martial Marbouty, Stéphane Duigou, Julien Mozziconacci, Olivier Espéli, Frédéric Boccard, and Romain Koszul. 2018. “Multiscale Structuring of the E. Coli Chromosome by Nucleoid-Associated and Condensin Proteins.” Cell 172 (4): 771–83.e18.

Magnusson, Klas E. G., Joakim Jalden, Penney M. Gilbert, and Helen M. Blau. 2015. “Global Linking of Cell Tracks Using the Viterbi Algorithm.” IEEE Transactions on Medical Imaging 34 (4): 911–29.

Marbouty, Martial, Antoine Le Gall, Diego I. Cattoni, Axel Cournac, Alan Koh, Jean-Bernard Fiche, Julien Mozziconacci, Heath Murray, Romain Koszul, and Marcelo Nollmann. 2015. “Condensin- and Replication-Mediated Bacterial Chromosome Folding and Origin Condensation Revealed by Hi-C and Super-Resolution Imaging.” Molecular Cell 59 (4): 588–602.

Masters, M. 1977. “The Frequency of P1 Transduction of the Genes of Escherichia Coli as a Function of Chromosomal Position: Preferential Transduction of the Origin of Replication.” Molecular & General Genetics: MGG 155 (2): 197–202.

Mirny, Leonid A. 2011. “The Fractal Globule as a Model of Chromatin Architecture in the Cell.” Chromosome Research: An International Journal on the Molecular, Supramolecular and Evolutionary Aspects of Chromosome Biology 19 (1): 37–51.

Nagano, Takashi, Yaniv Lubling, Csilla Várnai, Carmel Dudley, Wing Leung, Yael Baran, Netta Mendelson Cohen, Steven Wingett, Peter Fraser, and Amos Tanay. 2017. “Cell-Cycle Dynamics of Chromosomal Organization at Single-Cell Resolution.” Nature 547 (7661): 61–67.

Nielsen, Henrik J., Jesper R. Ottesen, Brenda Youngren, Stuart J. Austin, and Flemming G. Hansen. 2006. “The Escherichia Coli Chromosome Is Organized with the Left and Right Chromosome Arms in Separate Cell Halves.” Molecular Microbiology 62 (2): 331–38.

Niki, H., Y. Yamaichi, and S. Hiraga. 2000. “Dynamic Organization of Chromosomal DNA in Escherichia Coli.” Genes & Development 14 (2): 212–23.

Oluwadare, Oluwatosin, Max Highsmith, and Jianlin Cheng. 2019. “An Overview of Methods for Reconstructing 3-D Chromosome and Genome Structures from Hi-C Data.” Biological Procedures Online 21 (1): 7.

Price, Morgan N., Kelly M. Wetmore, Adam M. Deutschbauer, and Adam P. Arkin. 2016. “A Comparison of the Costs and Benefits of Bacterial Gene Expression.” PloS One 11 (10): e0164314.

Ramani, Vijay, Xinxian Deng, Ruolan Qiu, Kevin L. Gunderson, Frank J. Steemers, Christine M. Disteche, William S. Noble, Zhijun Duan, and Jay Shendure. 2017. “Massively Multiplex Single-Cell Hi-C.” Nature Methods 14 (3): 263–66.

Sadhir, Ismath, and Seán M. Murray. 2023. “Mid-Cell Migration of the Chromosomal Terminus Is Coupled to Origin Segregation in Escherichia Coli.” Nature Communications 14 (1): 7489.

Soares, Ruben R. G., Daniela A. García-Soriano, Jimmy Larsson, David Fange, Dvir Schirman, Marco Grillo, Anna Knöppel, et al. 2023. “Pooled Optical Screening in Bacteria Using Chromosomally Expressed Barcodes.” bioRxiv. 10.1101/2023.11.17.567382.

Thomason, Lynn C., Nina Costantino, and Donald L. Court. 2007. “E. Coli Genome Manipulation by P1 Transduction.” Et Al [Current Protocols in Molecular Biology] Chapter 1 (1): 1.17.1–1.17.8.

Umbarger, Mark A., Esteban Toro, Matthew A. Wright, Gregory J. Porreca, Davide Baù, Sun-Hae Hong, Michael J. Fero, et al. 2011. “The Three-Dimensional Architecture of a Bacterial Genome and Its Alteration by Genetic Perturbation.” Molecular Cell 44 (2): 252–64.

Valens, Michèle, Stéphanie Penaud, Michèle Rossignol, François Cornet, and Frédéric Boccard. 2004. “Macrodomain Organization of the Escherichia Coli Chromosome.” The EMBO Journal 23 (21): 4330–41.

Wallden, Mats, David Fange, Ebba Gregorsson Lundius, Özden Baltekin, and Johan Elf. 2016. “The Synchronization of Replication and Division Cycles in Individual E. Coli Cells.” Cell 166 (3): 729–39.

Wang, Xindan, Xun Liu, Christophe Possoz, and David J. Sherratt. 2006. “The Two Escherichia Coli Chromosome Arms Locate to Separate Cell Halves.” Genes & Development 20 (13): 1727–31.

Wasim, Abdul, Palash Bera, and Jagannath Mondal. 2023. “Development of a Data-Driven Integrative Model of a Bacterial Chromosome.” Journal of Chemical Theory and Computation, April. 10.1021/acs.jctc.3c00118.

Wasim, Abdul, Ankit Gupta, and Jagannath Mondal. 2021. “A Hi–C Data-Integrated Model Elucidates E. Coli Chromosome’s Multiscale Organization at Various Replication Stages.” Nucleic Acids Research 49 (6): 3077–91.

Yildirim, Asli, and Michael Feig. 2018. “High-Resolution 3D Models of Caulobacter Crescentus Chromosome Reveal Genome Structural Variability and Organization.” Nucleic Acids Research 46 (8): 3937–52.

Youngren, Brenda, Henrik Jörk Nielsen, Suckjoon Jun, and Stuart Austin. 2014. “The Multifork Escherichia Coli Chromosome Is a Self-Duplicating and Self-Segregating Thermodynamic Ring Polymer.” Genes & Development 28 (1): 71–84.

Zhang, Y., F. Buchholz, J. P. Muyrers, and A. F. Stewart. 1998. “A New Logic for DNA Engineering Using Recombination in Escherichia Coli.” Nature Genetics 20 (2): 123–28.

