## Supplemental figures and tables for "A dynamic 3D polymer model of the *Escherichia coli* chromosome driven by data from optical pooled screening"

### Supplemental Information

15 strains from the pool were not observed in enough growth channels using on-chip genotyping for reliable statistics (<10 channels) and were discarded from the analysis. 14 out of them were not found using sequencing as well, probably due to cloning biases or growth defects in these specific strains. One other strain was observed in the pool at the expected frequency using sequencing but was found only at 7 growth channels using on-chip genotyping, probably due to a low efficiency of this strain's barcode sequence.

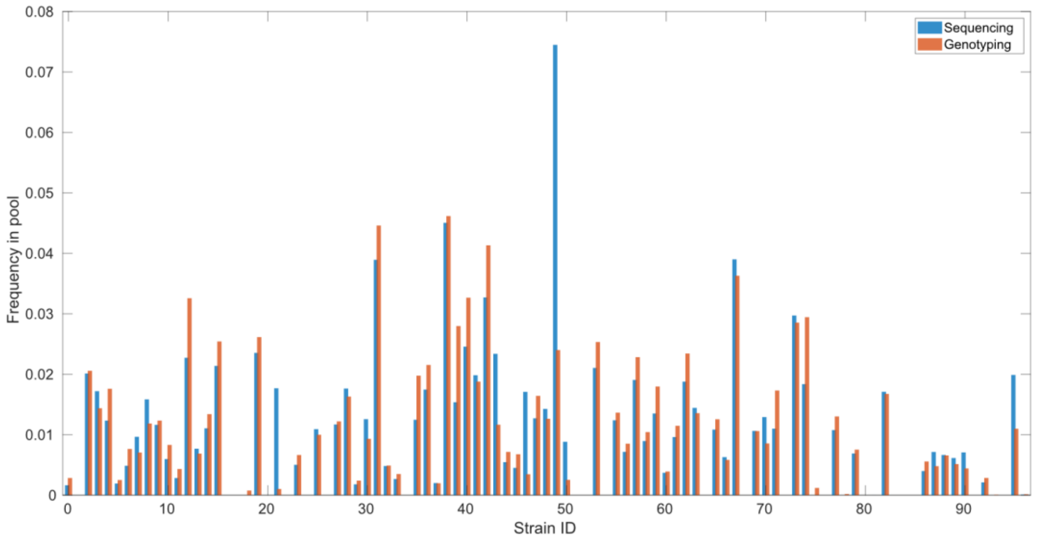

**Fig. S1. Comparing sequencing of the pooled strain library with genotyping**. Frequency of each strain in the library estimated with amplicon deep sequencing and genotyping.

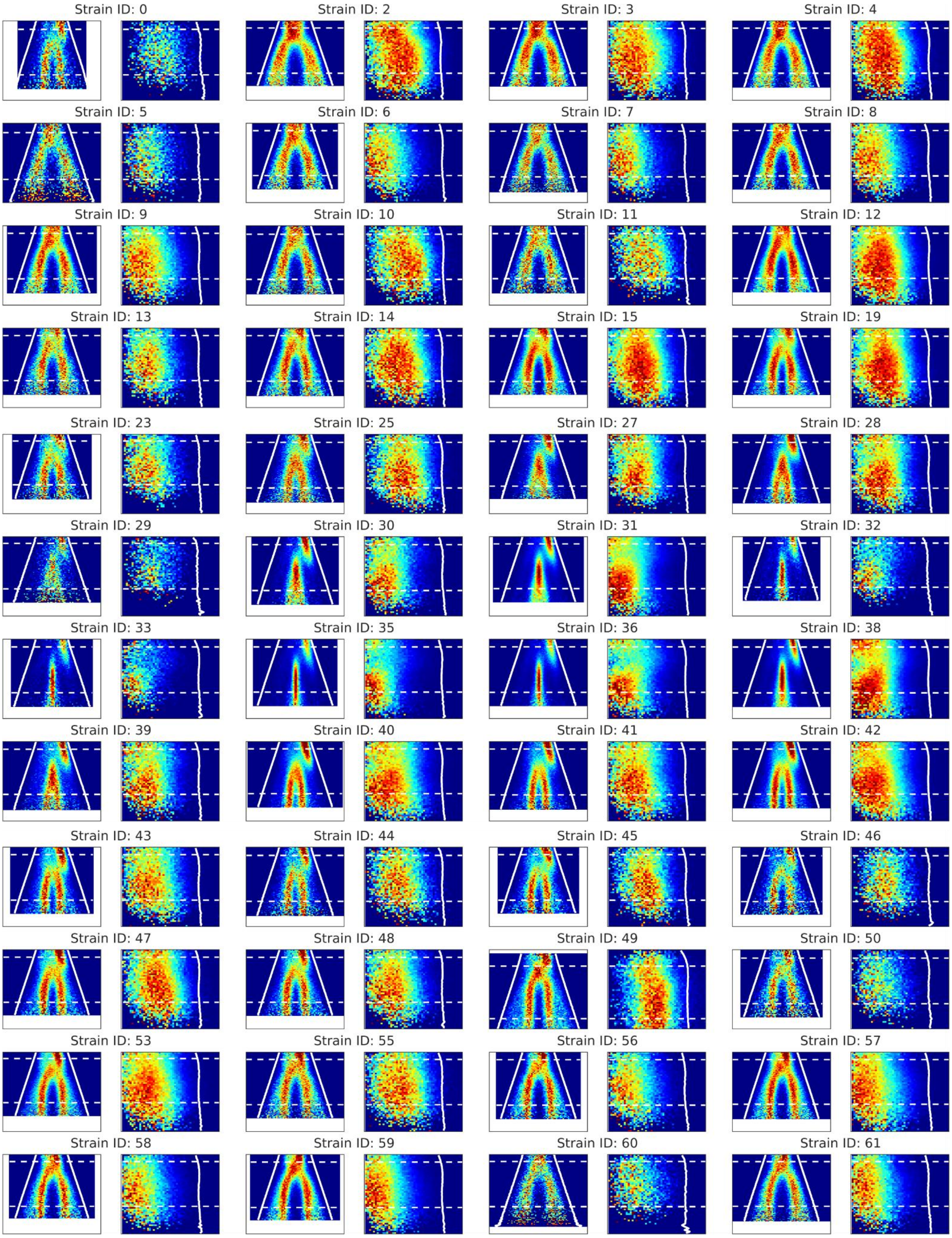
Strain ID: 0
Strain ID: 2
Strain ID: 3
Strain ID: 4
Strain ID: 5
Strain ID: 6
Strain ID: 7
Strain ID: 8
Strain ID: 9
Strain ID: 10
Strain ID: 11
Strain ID: 12
Strain ID: 13
Strain ID: 14
Strain ID: 15
Strain ID: 19
Strain ID: 23
Strain ID: 25
Strain ID: 27
Strain ID: 28
Strain ID: 29
Strain ID: 30
Strain ID: 31
Strain ID: 32
Strain ID: 33
Strain ID: 35
Strain ID: 36
Strain ID: 38
Strain ID: 39
Strain ID: 40
Strain ID: 41
Strain ID: 42
Strain ID: 43
Strain ID: 44
Strain ID: 45
Strain ID: 46
Strain ID: 47
Strain ID: 48
Strain ID: 49
Strain ID: 50
Strain ID: 53
Strain ID: 55
Strain ID: 56
Strain ID: 57
Strain ID: 58
Strain ID: 59
Strain ID: 60
Strain ID: 61

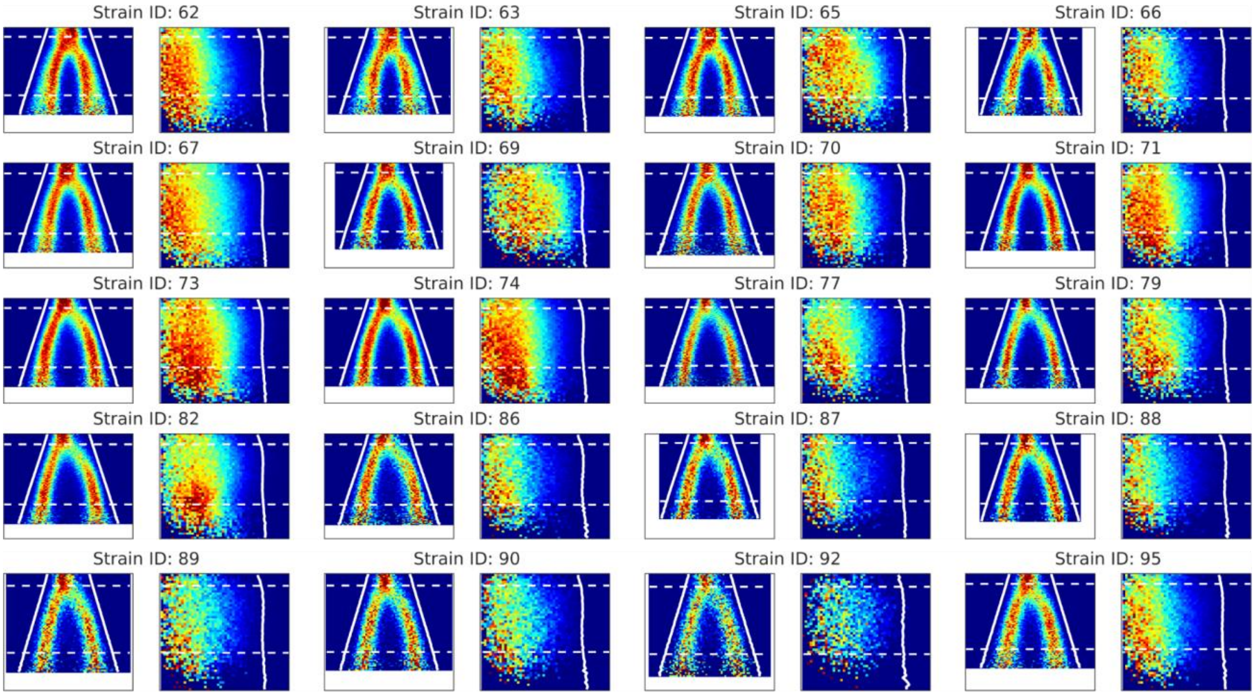

**Fig. S2**. Location distributions along the cell's long and radial axes as a function of cell area as in Fig. 2, but for all loci in the strain library. Strain 82 has the locus label closest to *oriC* (34 kb from *oriC* on the right chromosome arm) and strain 35 has the locus label in the middle of the *ter* region (47 kb from *dif*).

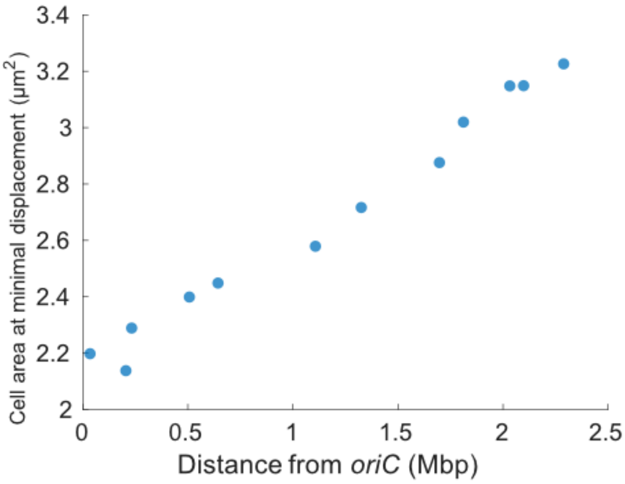

**Fig. S3**. Cell area at the average 1 s frame-to-frame displacement minimum from (Gras, Fange, and Elf 2024) for 12 chromosome loci in the strain library as function of distance from *oriC*.

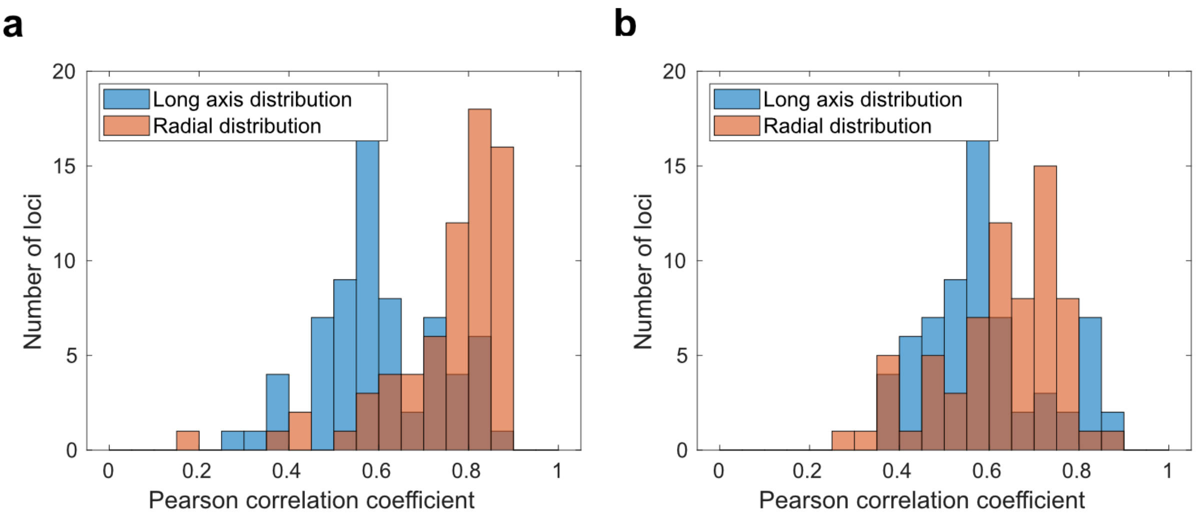

**Fig. S4. Pearson correlation coefficients for a randomized model.** Histograms of Pearson correlation coefficients between experimental and simulated long and radial axes coordinate distributions as in Fig. 4c and 4d, but for a polymer model with randomized location inputs.

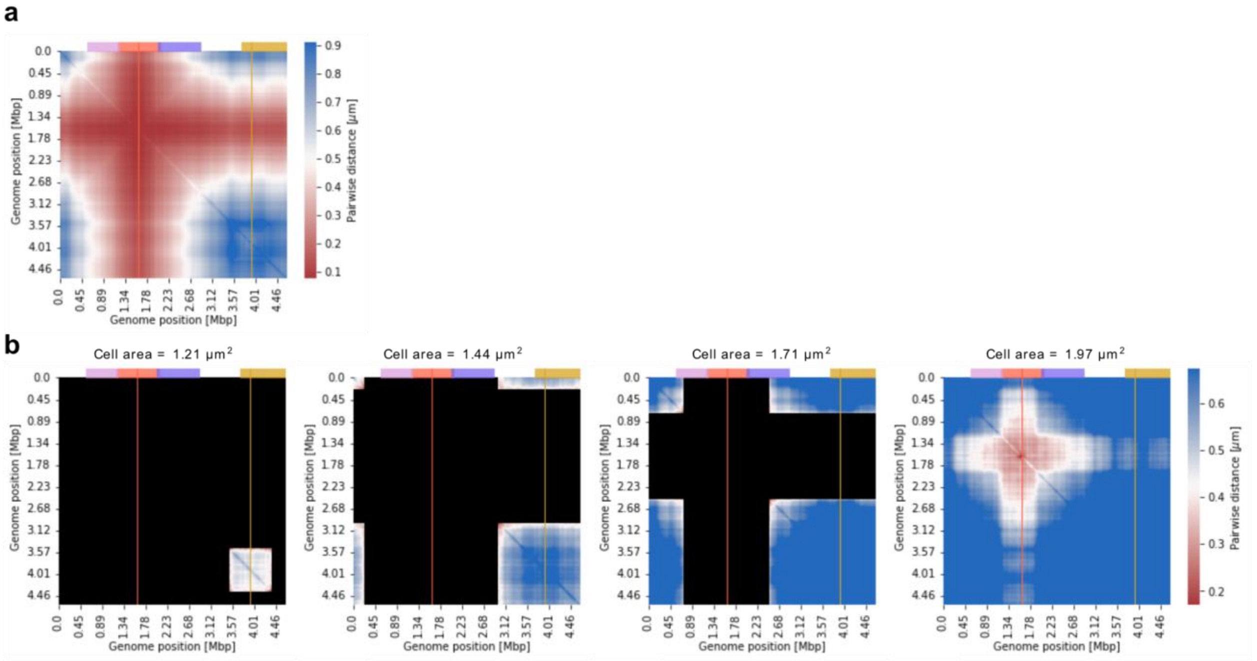

**Fig. S5. Dynamic interchromosomal distance maps generated with the polymer model.** Distance maps in **a,** and **b,** as in Fig. 6c and 6d, but for interchromosomal distances estimated between replicated loci in the polymer model. Each distance map in b, corresponds to a selected cell area bin.

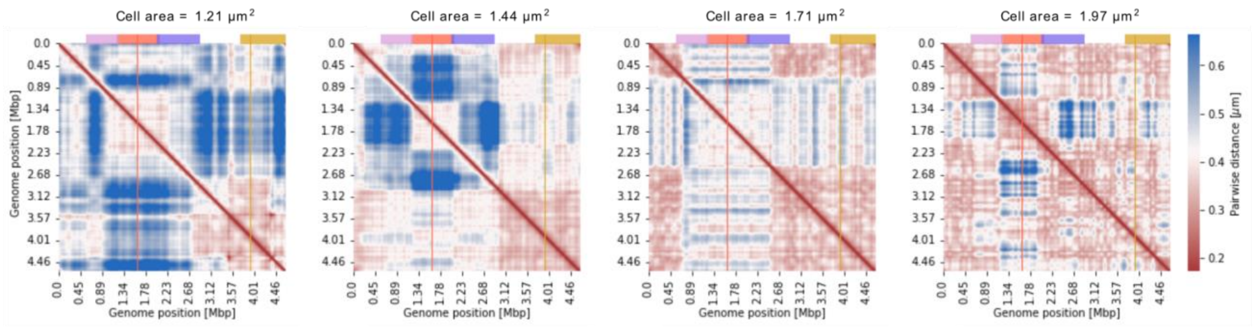

**Fig. S6. Chromosome distance maps generated using a segregating polymer with randomized**

**coordinate inputs.** As in the Fig. 6c, but for pairwise locus distances based on a polymer model with randomized loci coordinate inputs and that maintains chromosome segregation.

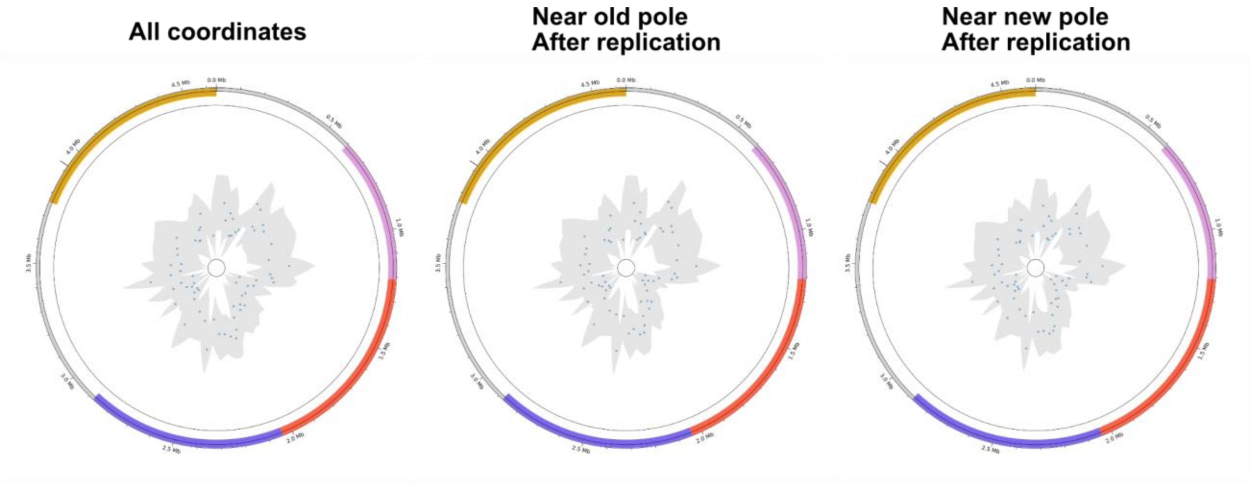

**Fig. S7**. Location distributions along the cell's radial axes as a function of cell area as in Fig. 4 for all loci in the strain library and for the replicated loci copies. For each locus label, the leftmost panel shows the radial location distribution and the panels labeled old pole and new pole show the radial location distribution of the loci copies closest to the old cell pole or the new cell pole. The cell area at which loci replicate was estimated using the replication fork progression modeled using Fig. S3.

In our previous study (Gras, Fange, and Elf 2024), we measured at what point during the cell cycle twelve specific loci replicated. Here, we used those results to infer replication progression as a function of genomic locus position (Fig. S3). This allowed us to compare the radial positioning of the replicated copies of the same locus for all of the library's loci following their replication and segregation by separating the copies based on their proximity to the old and new cell poles (Fig. S7). We did not find any significant difference in the radial location distributions between the loci before replication and either of its replicated copies, suggesting that the loci occupy similar intracellular positions radially before and after their replication.

**Table S1. Genotyping statistics for each experiment.** 1- After performing the enzymatic steps some channels were left with no cells' lysate (see methods for how we identify these channels) 2- Channels that included cells' lysate but produced no fluorescence signal during the rounds of sequential probing. 3 - Channels that fluoresced in more than one color at a given sequential probing round. 4 - Channels where the readout sequence of detection probes was more than one symbol away from any barcode in the library and could thus not be error corrected. 5 - Number of channels that produced a valid barcode signal that was successfully assigned to a library barcode either directly or using error correction. The percentage in the brackets represents the percentage of successfully decoded channels among channels that include cells' lysate.

| Exp. id | Nr. of growth channels | Nr. of empty channels[1] | Channels with no signal[2] | Channels with double signal[3] | Unassigned decoded channels[4] | Successfully decoded channels[5] |
|---|---|---|---|---|---|---|
| EXP-23-CA3093 | 2000 | 95 | 149 | 209 | 51 | 1496 [78.5%] |
| EXP-23-CA3099 | 1760 | 8 | 151 | 224 | 59 | 1318 [75.23%] |
| EXP-24-CB4759 | 2000 | 12 | 212 | 277 | 141 | 1358 [68.31%] |
| EXP-24-CB4763 | 4000 | 54 | 311 | 219 | 192 | 3220 [81.6%] |

**Table S2. List of oligonucleotides.**

| Name | Notes | Sequence |
|---|---|---|
| prDS147 | Amplifying R6K vector for plasmid EL3395 construction | atccagcagttcaacctg |
| prDS148 | Amplifying R6K vector for plasmid EL3395 construction | gggagaccagaaacaaaaaaag |
| prDS149 | Ampliying barcode expression 3' for plasmid EL3395 construction | ttttttgtttctggtctccctcaaccaataatagtctgaatg |
| prDS150 | Amplifying barcode expression 3' for plasmid EL3395 construction | tgcgcttgcgcaaaaaacccctcaagac |
| prDS151 | Amplifying kanR for plasmid EL3395 construction | gggttttttgcgcaagcgcaaagagaaag |
| prDS152 | Amplifying kanR for plasmid EL3395 construction | gccagggccatcagaagaactcgtcaagaag |
| prDS153 | Amplifying malOx12 array for plasmid EL3395 construction | gttcttctgatggccctggcatcacctc |
| prDS154 | Amplifying malOx12 array for plasmid EL3395 construction | aacaggttgaactgctggatcacttacttggttcaacctcagagag |
| prDS292 | Amplifying barcode region for amplicon sequencing | ACACTCTTTCCCTACACGACGCTCTTCCGATCTtaatacgactcactatagggag |
| prDS293 | Amplifying barcode region for amplicon sequencing | AGACGTGTGCTCTTCCGATCcaaaaaacccctcaagacc |

**Table S3. Library strains and associated oligonucleotide sequences.**
***** Label insertion sites are indicated based on *E. coli* str. K-12 substr. MG1655 complete genome (U00096.3).

**Table S4. Reaction mixes used for the genotyping protocol.**

| 1. Lysozyme reaction mix | Stock Conc. | Final Conc. | Volume (µL) |
|---|---|---|---|
| Lysozyme (Thermo scientific) | 50 mg/ml. | 250 µg/ml | |
| PBS-Tween 0.01% | | | |
| **2. BSA solution** | **Stock Conc.** | **Final Conc.** | **Volume (µL)** |
| BSA (Thermo scientific) | 10% | 1% | |
| PBS-Tween 0.01% | | | |
| **3. Zombie transcription mixture** | **Stock Conc.** | **Final Conc.** | **Volume (µL)** |
| Transcription buffer (Thermo scientific) | 5x | 1x | 16 |
| NTPs | 25 mM | 2 mM | 6.4 |
| MgCl2 | 25 mM | 6 mM | 19.2 |
| Tween 20 (Promega) | 1% (v/v) | 0.1% (v/v) | 8 |
| Glycerol | 50% | 5% | 8 |
| T7 RNA Polymerase (Thermo scientific) | 20 U/µl | 2 U/µl | 8 |
| Riboprotect (Qiagen gdansk) | 40 U/µl | 1 U/µl | 2 |
| H2O mQ | - | - | 12.4 |
| **4. PLP Hybridization mixture** | **Stock Conc.** | **Final Conc.** | **Volume (µL)** |
| 20x SSC | 20x | 2x | 5 |
| Ethylene Carbonate (Merck) | 100% | 5% | 2.5 |
| MgCl2 | 50 mM | 15 mM | 15 |
| Tween 20 | 1% (v/v) | 0.1% (v/v) | 5 |
| Padlock probe library (each probe) | 0.41 µM | 175 nM | 21.25 |
| Riboprotect (Qiagen gdansk) | 40 U/µl | 1 U/µl | 1.25 |
| **5. SplintR ligation mixture** | **Stock Conc.** | **Final Conc.** | **Volume (µL)** |
| SplinR Buffer (NEB) | 10x | 1x | 6 |
| Glycerol | 50% | 5% | 6 |

| | | | |
|---|---|---|---|
| Tween 20 | 1% (v/v) | 0.1% (v/v) | 6 |
| Riboprotect (Qiagen gdansk) | 40 U/µl | 1 U/µl | 1.5 |
| SplintR Ligase (NEB) | 25 U/µL | 0.5 U/µL | 1.2 |
| H2O mQ | - | - | 39.3 |
| **6. RCA primer hybridization mixture** | **Stock Conc.** | **Final Conc.** | **Volume (µL)** |
| 20x SSC | 20x | 2x | 5 |
| Ethylene Carbonate (Merck) | 100% | 5% | 2.5 |
| MgCl2 | 50 mM | 15 mM | 15 |
| Tween 20 | 1% (v/v) | 0.1% (v/v) | 5 |
| Glycerol | 50% | 1.25% | 1.25 |
| RCA primer **GGCTCCACTAAATAGACGCA** | 10 µM | 1 µM | 5 |
| H2O mQ | - | - | 16.25 |
| **7. RCA mixture** | **Stock Conc.** | **Final Conc.** | **Volume (µL)** |
| dNTPs | 2.5 mM | 250 uM | 9 |
| phi 29 buffer * | 10x | 1 x | 9 |
| Glycerol | 50% | 5% | 9 |
| BSA | 20 µg/µl | 0.2 µg/µl | 0.9 |
| phi 29 polymerase * | 10 U/uL | 1 U/uL | 9 |
| H2O mQ | - | - | 53.1 |
| **8. Labeling mixture** | **Stock Conc.** | **Final Conc.** | **Volume (µL)** |
| Detection oligos (IDT) | 1 µM each | 100 nM | 6 |
| 4x SSC 40% Formamide | 2x | 1x | 30 |
| L-probe pool | 1 µM total | 0.1 µM total | 6 |
| H2O mQ | - | - | 18 |
| **9. Probe stripping mixture** | **Stock Conc.** | **Final Conc.** | **Volume (µL)** |
| Formamide | 100% | 90% | 900 |
| PBS-Tween | 1x | 0.1x | 100 |

* Wild-type phi29 DNA polymerase was transformed into E. coli BL21 (DE3) T1R pRARE2 cells. The cells were cultivated in Terrific Broth (TB) medium. Protein expression was induced with IPTG and the protein was purified by immobilized metal-ion chromatography (IMAC), followed by size exclusion chromatography (SEC).
† The phi29 reaction buffer at 10x concentration is: 500 mM Tris-HCl (pH 8.3), 100 mM MgCl2 & 100 mM (NH4)2SO4

**Table S5. Genotyping code and list of genotyping oligo pools**
This table describes the probe pools used for the genotyping experiments. The probe pools were mixed for a bigger library pool that included 97 strains. Probes that are associated with a library strain have a "strainID" indicated in the relevant column. Probes with no "strainID" exist in the probe pool but no corresponding strain exists in the strain library.
