## Supplementary figures and images for "A dynamic 3D polymer model of the *Escherichia coli* chromosome driven by data from optical pooled screening"

### Supplemental movie 1

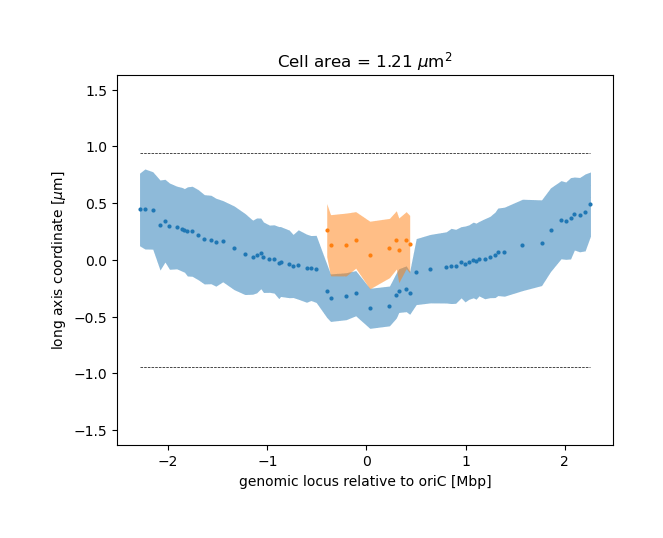

### Supplemental movie 2

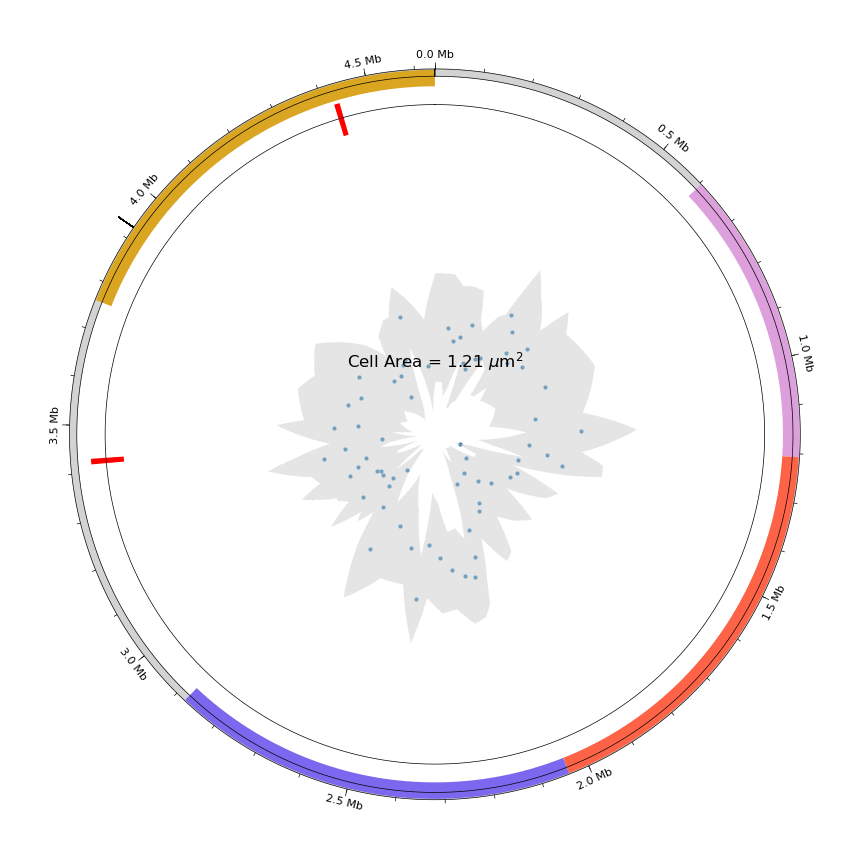

### Supplemental movie 4

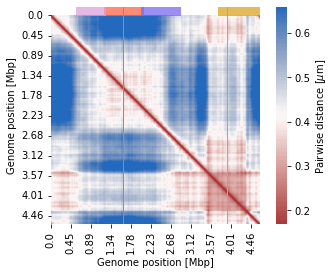

### Supplemental movie 5

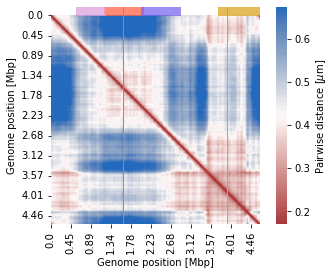

### Supplemental movie 6

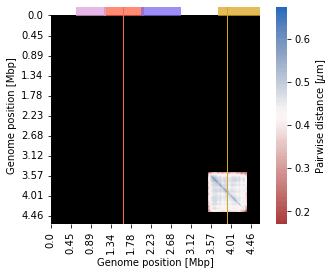
